# *Coxiella burnetii* type IVB secretion system modulates TLR3/TRIF-dependent NF-κB and IRF responses during infection

**DOI:** 10.64898/2026.09.23.753975

**Authors:** Saugata Mahapatra, Aryashree Arunima, Anna Busbee, Jennifer E. Dumaine, Lauren Stranahan, Ricardo A. Zeledon, Bobby Dow, Samantha Butler, Sabrina Clark, A. Phillip West, Erin J. van Schaik, James E. Samuel

## Abstract

*Coxiella burnetii* (Cb), the causative agent of Q fever, replicates within host macrophages by modulating innate immune responses through its type IVB secretion system (T4SS). Host cells sense pathogens via Pattern recognition receptors (PRRs), including Toll-like receptors (TLRs), to recognize pathogen-associated molecular patterns (PAMPs) and initiate signaling pathways that drive pro-inflammatory cytokine and interferon responses. Toll-like receptor 3 (TLR3) triggers the nuclear translocation of nuclear factor kappa-light-chain-enhancer of activated B cells (NF-κB) and Interferon regulatory factors (IRFs) resulting in pro-inflammatory cytokine production and type I interferon (IFN-I) induction. Here we demonstrate that Cb requires T4SS to suppress TLR3-induced NF-κB and IRF transcriptional responses. Furthermore, RNA purified from both virulent and avirulent Cb is sufficient to activate TLR3, identifying pathogen RNA as a relevant PAMP in this context. Using a pulmonary infection model with virulent Cb, we found that TLR3/ Toll-Interleukin-1 Receptor Domain-Containing Adapter Protein Inducing Interferon Beta (TRIF)-dependent innate immune pathway inhibits cachexia. In contrast, signaling through the interferon-α/β receptor (IFNAR) restricts bacterial dissemination, indicating that these pathways play distinct but complementary roles in host defense. Mechanistically, Cb suppresses TLR3/TRIF signaling in a T4SS-dependent manner by preventing host TLR3 recruitment to the Coxiella-containing vacuole (CCV) and disrupting TRIF–TNF Receptor-Associated Factor 6 (TRAF6) interactions, thereby selectively inhibiting impairing NF-κB activation. Consistent with this, we identify five T4SS effector proteins that attenuate TLR3-induced NF-κB signaling, including CBU1292, which suppresses both NF-κB and IRF-dependent responses downstream of TLR3.

**Author summary:** Pathogenic bacteria have evolved mechanisms to evade or manipulate host immune defenses. *Coxiella burnetii* (Cb), the causative agent of Q fever, replicates within immune cells and interferes with innate immune signaling pathways. We found that its type IVB secretion system (T4SS) express effectors that inhibit immune responses triggered by Toll-like receptor 3 (TLR3), a sensor that detects microbial RNA. Specifically, Cb suppresses signaling through the adaptor protein TRIF, thereby reducing activation of key immune regulators, NF-κB and interferon regulatory factors (IRFs), which are required for inflammatory and antiviral responses. Using a mouse model of lung infection, we found that disruption of TLR3/TRIF signaling and type I interferon receptor (IFNAR) signaling leads to different disease outcomes, indicating that these pathways play distinct roles in controlling infection and disease severity. We also identified several Cb-secreted effectors that interfere with TLR3-mediated immune activation, including one that inhibits both NF-κB and interferon responses. Together, our findings reveal how Cb evades immune detection by targeting a critical RNA-sensing pathway and highlight distinct host defense mechanisms that limit bacterial infection and disease.

## Introduction

*Coxiella burnetii* (Cb) is a Gram-negative obligate intracellular bacterium that infects alveolar macrophages and causes Q fever in humans (1, 2). This environmentally stable pathogen exhibits an exceptionally low infectious dose and is acquired by inhalation of contaminated particles, rendering it a CDC select agent (2, 3). After phagocytosis and acidification of the endosome, Cb deploys its type IVB secretion system (T4SS) to manipulate host cellular pathways, promoting heterotypic fusion that creates a single membrane-bound replicative compartment termed the *Coxiella*-containing vacuole (CCV) (1, 4–7). Most *in vitro* studies utilize the phase II avirulent variant NMII (Cb Nine Mile phase II RSA439), which contains a large deletion in the lipopolysaccharide (LPS) O-antigen biosynthesis genes that exempts it from select agent restrictions, and allows BSL2 handling (4–9). However, due to NMII’s high attenuation, its incapable of causing disease in immunocompetent animal models including mice and guinea pigs, which creates a disconnect between *in vitro* findings and the virulence mechanisms required for Cb pathogenesis *in vivo* (10, 11).

A hallmark of Cb infection is its ability to suppress host innate immune responses while establishing prolonged intracellular replication (12). This immune evasion is exemplified by Cb’s capacity to replicate within mouse bone marrow-derived macrophages (BMDM) for multiple days with minimal innate immune activation or host cell death, even as the CCV occupies substantial cell cytoplasmic space (13, 14). Studies using knockout mice demonstrate that deletion of key Pattern Recognition Receptors (PRRs), including Toll-like receptors (TLRs) and their downstream adapter proteins, creates more permissive environments for Cb replication. Enhanced bacterial growth in TLR2^−^/^−^, TLR4^−^/^−^, MyD88^−^/^−^, or TRIF^−^/^−^ BMDMs provides evidence that these TLR-specific signaling pathways are likely detrimental to Cb survival (15–18).

TLR activation triggers signaling cascades that induce nuclear factor kappa-light-chain-enhancer of activated B cells (NF-κB)-dependent pro-inflammatory responses and/or Interferon Regulatory Factor (IRF)-dependent type-I interferon (IFN-I) production (18–20). These IFN-I responses signal through the interferon-α/β receptor (IFNAR) to drive a broader antimicrobial immune response (18, 19). *In vivo* studies reveal complex tissue-specific effects: while IFNAR^−^/^−^ mice show reduced cytokine levels in bronchoalveolar lavage fluid to Cb infection (18), Cb replication in the lung is not enhanced, suggesting that IFN-I signaling may promote bacterial replication in peripheral tissues while restricting pulmonary infection (18, 19). Supporting this observation, TLR2^−^/^−^ mice exhibit increased permissiveness to pulmonary infection with virulent Cb Nine Mile phase I (NMI), and both TLR2^−^/^−^ and MyD88^−^/^−^ mice support enhanced NMII colonization following intraperitoneal infection (17, 21, 22). Additionally, IFNAR^−^/^−^ mice infected with NMI show reduced weight loss compared to wildtype mice, while intratracheal recombinant IFN-α treatment in wildtype mice prevents Cb dissemination to the spleen (18).

These findings indicate that NF-κB activation via the TLR2/MyD88 pathway, together with IFN-I signaling downstream of IFNAR, contributes to restricting Cb replication *in vivo*. This aligns with reports demonstrating that pro-inflammatory NF-κB-driven responses, including TNF-α production, play important roles in suppressing Cb growth both *in vitro* and *in vivo*. Moreover, its established that Cb dampens NF-κB activation through T4SS-dependent mechanisms (12, 23, 24). Both IFN-α and IFN-γ can restrict Cb growth in macrophages, with functional crosstalk between these pathways and TNF-α-mediated bacterial restriction (15, 25). The discovery of Cb T4SS effectors that modulate NF-κB and IFN-I responses, including NopA, CBU1314, CBU1639, CinF, EmcA, and EmcB, underscores the importance of these pathways during infection, and suggests that additional effectors likely act downstream of TLRs and/or IFNAR to promote infection (13, 25–29).

Despite this progress, the upstream ligands driving these pathways remain poorly defined. Cb LPS is largely antagonistic to TLR4 due to its atypical tetra-acylated lipid A structure and induces weak TNF-α release in mouse BMDM. Importantly, lipid A structure is conserved between virulent NMI and avirulent NMII, indicating minimal, phase-independent TLR4 signaling (24, 30). While TLR2 is important for Cb infection both *in vitro* and *in vivo*, relevant ligands remain unknown despite evidence that Cb stimulates TLR2 and TLR1/2 signaling in human PBMC (31). These observations suggest that additional Cb pathogen-associated molecular patterns (PAMPs) and T4SS effectors remain to be identified that regulate TLR or IFN-I-dependent pathways. Mechanistically, TLR signaling occurs through cytoplasmic TIR domains that interact with either MyD88 (TLR1/2, TLR2, TLR4, TLR5, TLR2/6, TLR7, TLR9) or TRIF (TLR3 and endosomal TLR4). TLR/MyD88 signaling primarily drives NF-κB-dependent inflammatory gene expression, whereas TLR/TRIF signaling predominantly induces IFN-I production through IRF3, while also activating NF-κB. (20, 32). The recent discovery of EmcB, which removes K63-linked ubiquitin chains to inhibit RIG-I, suggests that Cb may generate dsRNA capable of activating RIG-I and potentially TLR3 (25). Consistent with this, the observation of formalin-fixed vaccine material eliciting a non-specific antiviral response supports the presence of a Cb-derived PAMP that can trigger IFN-I pathways which are likely actively inhibited during infection (33, 34).

In this study, we investigated whether Cb utilizes its T4SS to modulate host TLR signaling pathways and their contribution to infection control. We report for the first time that Cb employs its T4SS to suppress the host TLR3/TRIF-induced innate immune signaling pathway, thereby inhibiting both NF-κB and IFN-I responses. We demonstrate that RNA purified from both virulent NMI and avirulent NMII strains activates TLR3 signaling. Employing a mouse pulmonary infection model, we demonstrate that both the TLR3/TRIF signaling axis and IFNAR signaling contribute to distinct aspects of host defense. Mechanistically, Cb disrupts TRIF-TRAF6 interactions in a T4SS-dependent manner, suppressing NF-κB signaling. Finally, screening a Cb Himar1 transposon mutant library containing 32 T4SS effector candidates identified several effectors contributing to NF-κB suppression, including CBU1292, which inhibits both NF-κB and IRF-I signaling (35, 36). Collectively, these findings suggest that Cb deploys its effector repertoire to disrupt the host’s key innate immune signaling networks, specifically targeting TLR3/TRIF-induced antimicrobial sensing through multiple mechanisms.

## Results

### Cb employs the T4SS to suppress host TLR3-induced NF-κB-signaling during infection

To determine whether Cb modulates TLR-induced NF-κB activation through its T4SS, we employed THP1-Lucia™ NF-κB Cells (InvivoGen), which harbor an NF-κB-inducible secreted Lucia luciferase reporter gene (NF-κB-Lucia). NF-κB activity was quantified by measuring secreted luciferase activity in the infected culture supernatant. These cells were infected with either wild-type Cb NMII or a T4SS-deficient NMII Δ*dotA* mutant (DotA). Multiplicity of infection (MOI) was optimized using immunofluorescence microscopy to ensure close to 100% infection rates for both strains, eliminating differential infection as a confounding variable (S1 Fig). During 24-hour infection monitoring without TLR agonist stimulation, significant differences were observed in NF-κB-Lucia activation between DotA and NMII-infected cells beginning at 6 hours post-infection (hpi) through 24 hpi (one-way ANOVA with Tukey’s test, p < 0.05; Fig 1A). These findings indicate that the Cb T4SS actively dampens NF-κB signaling and are consistent with previous studies demonstrating T4SS-dependent suppression of NF-κB signaling (37).

**Fig. 1.**
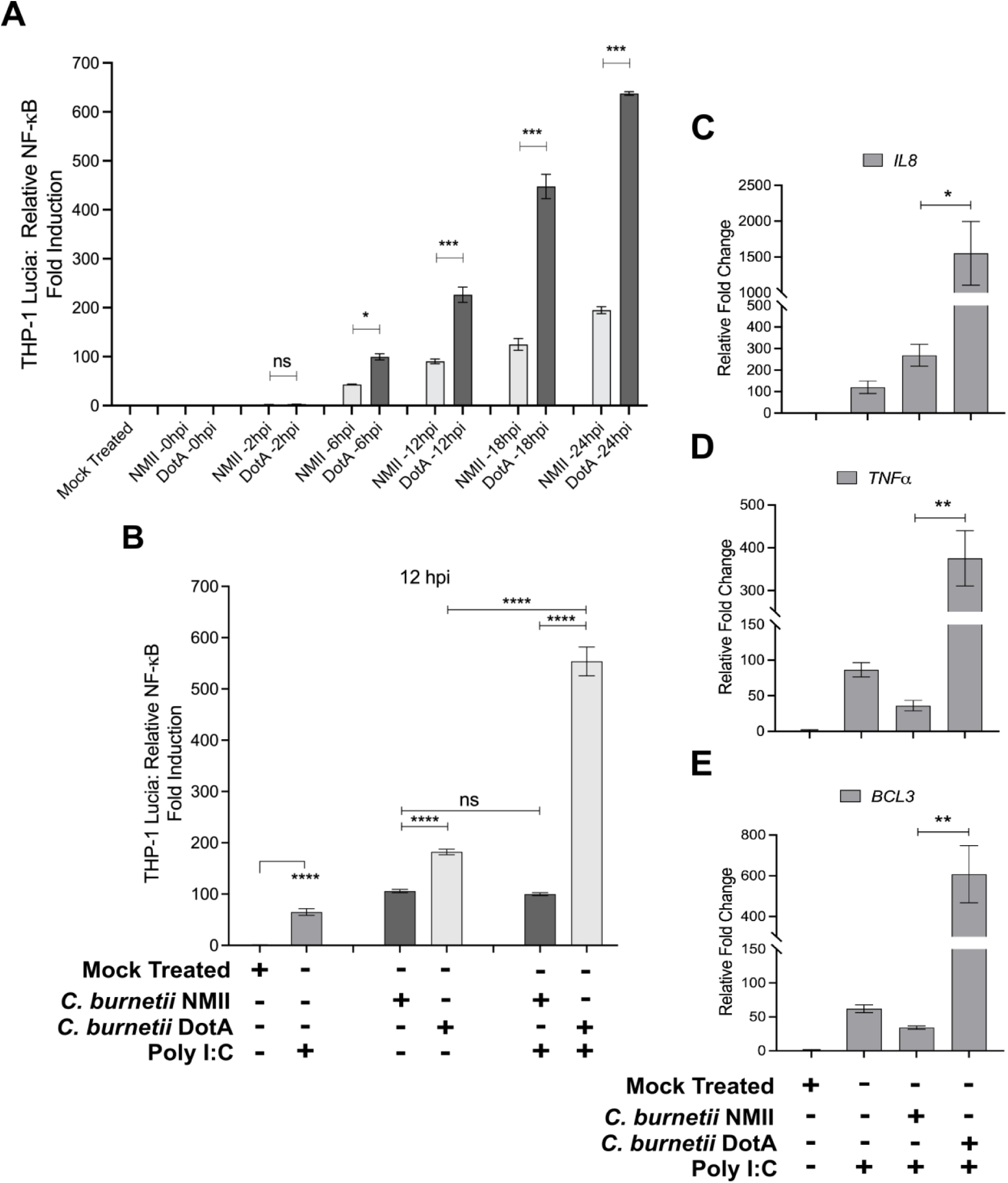
Cb suppresses the TLR3-induced NF-κB-activation in a T4SS-dependent manner. Luciferase assay measuring the NF-kB-Lucia activity in NMII vs DotA infected cells. A 24-hour time-course experiment was conducted with THP-1 Lucia cells that were either mock treated or infected with either NMII or DotA at an MOI of 25. Luminescence data was collected at the indicated time points. Data are presented as mean ± SEM from three independent experiments, with statistical significance (p-value < 0.05) determined for NMII vs. DotA at the indicated time points using ordinary one-way ANOVA with Tukey’s multiple comparison test. (B) Bioluminescent assay measuring the NF-κB-Lucia activity at 12 hpi in a THP-1 Lucia infection model to test the ability of wild-type NMII vs the DotA mutant to suppress TLR3-induced NF-κB activation. Cells were infected with NMII or DotA at an MOI of 25, followed by poly I:C stimulation (20 µg/ml) at 2 hpi. NF-κB-Lucia activation levels remained consistent (∼100-fold) in both unstimulated and poly I:C stimulated NMII-infected cells. In contrast, poly I:C stimulation in DotA-infected cells significantly increased NF-κB-Lucia activation from ∼200 to ∼550-fold, highlighting the T4SS-dependent suppression of TLR3-induced NF-κB activation by wild-type NMII. Data is represented as mean ± SEM of three independent experiments. Statistical comparisons were performed using two-way ANOVA with Tukey’s multiple comparison test (p-value < 0.05) for the following groups: Mock treated vs. 20 µg/ml poly I:C treated cells, NMII (poly I:C 0, 20 µg/ml) vs. DotA (poly I:C 0, 20 µg/ml), NMII (poly I:C 0 µg/ml) vs. NMII (poly I:C 20 µg/ml), and DotA (poly I:C 0 µg/ml) vs. DotA (poly I:C 20 µg/ml). (C-E) In wild-type THP-1 cell infections, RT-qPCR analysis of selected NF-κB-regulated host genes (*IL8*, *TNFα*, and *BCL3*) showed fold differences in mRNA levels relative to the *β-*actin control gene. THP-1 cells were infected with either wild-type NMII or the DotA mutant (MOI 25) for 2 hours, followed by stimulation with 20 µg/ml poly I:C for an additional 10 hours. Controls included THP-1 cells subjected to either mock treatment or poly I:C treatment, as indicated. Total RNA was harvested for RT-qPCR at 12 hpi. The x-axis represents the conditions under which gene expression was measured. The data represent the mean of three independent experiments, and error bars denote the standard error of the mean (SEM) with *p*-value calculated using two-way ANOVA with Tukey’s multiple comparisons test. (C) Relative fold changes in *IL8* mRNA levels, *p*-value < 0.05 for NMII + poly I:C (20 µg/ml) vs. DotA + poly I:C (20 µg/ml). (D) Relative fold changes in *TNFα* mRNA levels, *p*-value < 0.05 for NMII + poly I:C (20 µg/ml) vs. DotA + poly I:C (20 µg/ml). (E) Relative fold changes in *BCL3* mRNA levels, *p*-value < 0.05 for NMII + poly I:C (20 µg/ml) vs. DotA + poly I:C (20 µg/ml).

We next investigated whether Cb selectively disrupts specific TLR-induced NF-κB activation pathways. NMII or DotA infected THP1-Lucia cells were stimulated with TLR-specific agonists (S1 Table) at 2 hpi and analyzed for NF-κB-Lucia activity at 12 hpi. Distinct TLR pathways were selectively activated using PAM3CSK4 (for TLR2; S2 Fig), *E. coli* LPS (for TLR4; S3 Fig), poly I:C (for TLR3; S4 Fig), and CL075 (for TLR7/8; S5 Fig). Among these pathways, NMII effectively suppressed TLR3-induced NF-κB signaling in a T4SS-dependent manner. At 20 µg/ml poly I:C (Fig 1B; S4 Fig), NMII significantly suppressed TLR3-induced NF-κB activation (two-way ANOVA with Tukey’s test, p < 0.05). While NF-κB-Lucia activation in NMII-infected cells remained unaffected by poly I:C stimulation, DotA-infected cells exhibited a 3-fold increase in NF-κB-Lucia levels compared to unstimulated controls (two-way ANOVA with Tukey’s test, p < 0.05). These findings indicate that Cb preferentially suppresses TRIF-dependent over MyD88-dependent TLR signaling.

To validate these observations in parental THP-1 cells, we next examined the transcriptional induction of NF-κB-responsive genes following TLR stimulation. Three NF-κB-regulated genes (*IL8*, *TNFα*, and *BCL3*) were analyzed via RT-qPCR. TheTHP-1 cells were infected with either NMII or DotA and treated with poly I:C (TLR3 agonist) or *E. coli LPS* (TLR4 agonist) at 2 hpi. At 12 hpi, poly I:C stimulation (20 µg/ml) induced significantly higher mRNA expression of *IL8* (Fig 1C), *TNFα* (Fig 1D), and *BCL3* (Fig 1E) in DotA-infected compared to NMII-infected cells (two-way ANOVA with Tukey’s test, p < 0.05). Similar results were observed with 2 µg/ml poly I:C (S6A-C Fig). Conversely, TLR4 stimulation with *E. coli* LPS elevated *IL8*, *TNFα*, and *BCL3* mRNA levels comparably in both NMII- and DotA-infected cells (S7A-C, S8A-C Fig). These results further confirm that Cb T4SS selectively modulates TRIF-dependent TLR3 signaling without affecting MyD88-dependent TLR4 pathways.

### Cb T4SS suppresses TLR3-induced NF-κB and IRF signaling while Cb RNA functions as a PAMP

Our initial findings demonstrated that Cb NMII modulates NF-κB signaling through the TLR3–TRIF axis. Because TLR3 signaling is known to activate IFN-I responses, we next sought to determine whether Cb similarly suppresses IRF signaling in a T4SS-dependent manner. Since poly I:C can activate both TLR3 and RIG-I-like receptor pathways (38, 39), we employed a pharmacological TLR3 inhibitor (thiophene-carboxamidopropionate) to determine the specificity of TLR3-dependent signaling. We additionally hypothesized that Cb RNA functions as a PAMP engaging TLR3–TRIF signaling and that Cb suppresses both NF-κB and IRF responses through T4SS-dependent mechanisms. To test this, THP-1 Dual™ hTLR3 (InvivoGen) reporter cells were infected with either Cb NMII or DotA mutant with or without TLR3 inhibitor treatment. NF-κB and IRF activities were quantified at 24hpi using NF-κB-SEAP and IRF-luciferase assays, respectively. Cb NMII infection markedly suppressed both NF-κB and IRF responses relative to DotA infection, showing ∼5-fold and ∼13-fold reductions, respectively (Fig 2A and 2B). Interestingly, TLR3 inhibition only partially reduced the signaling in DotA-infected cells, suggesting that activation of additional PRRs could contribute to the activation of these pathways during infection.

**Fig 2.**
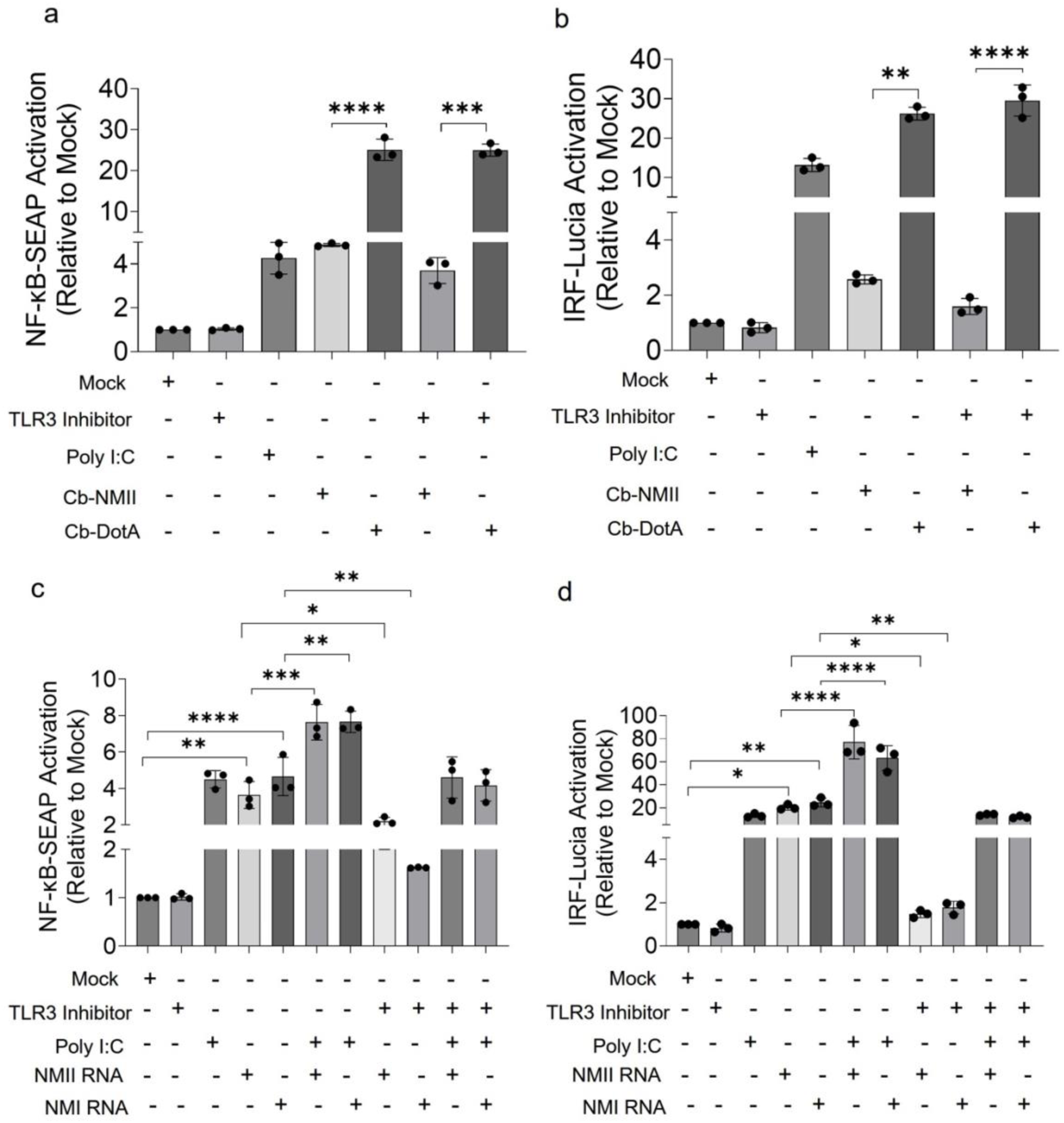
Cb suppresses NF-κB and IRF signaling via TLR3 in a T4SS-dependent manner. (a-d) THP-1-Dual-hTLR3 cells were infected with wild-type Cb NMII or the T4SS-deficient mutant DotA at an MOI of 25 for 3 h,or treated with RNA (1 µg/well) purified from virulent NMI or avirulent NMII. Cells were then stimulated with the TLR3 agonist poly(I:C) (20 μg/ mL) with or without the TLR3 inhibitor thiophene-carboxamidopropionate (10 μg/ mL), with standalone controls as indicated. NF-κB and IRF activation were quantified in cell culture supernatants by Lucia luciferase and SEAP assays, respectively. Activation is expressed as fold change relative to mock-treated cells (set to 1). Data is expressed as mean ± SD from three independent experiments, each performed in technical pentaplicates (*n* = 5 per condition). Statistical analysis: one-way ANOVA followed by Tukey’s or Dunnett’s multiple-comparison test, as appropriate. Significance: *, *p* < 0.05; **, *p* < 0.01; ***, *p* < 0.001; ****, *p* < 0.0001; ns, not significant, *p* ≥ 0.05.

Next, to determine whether Cb RNA could directly activate these pathways, DNA-free RNA purified from both virulent NMI and avirulent NMII strains (1 μg each) was used to stimulate THP-1 Dual™ hTLR3 cells alongside poly I:C and TLR3 inhibitor controls. Both NMI Cb and NMII Cb RNAs induced robust NF-κB and IRF activation in a TLR3-dependent manner, with stronger induction observed for IRF signaling. Maximal activation was observed in RNA- and poly I:C–treated cells (∼7.6-fold increase), whereas TLR3 inhibitor treatment reduced signaling by ∼2-fold (Fig 2C and 2D). These results suggest that Cb RNA functions as a TLR3 ligand capable of activating both NF-κB and IRF-I pathways.

To confirm TRIF as the downstream adaptor mediating this response, we analyzed NF-κB-SEAP and IRF-luciferase activities in TRIF^+^/^+^ and TRIF^−^/^−^ THP-1 Dual™ reporter (InvivoGen) cells infected with either Cb NMII or DotA or treated with bacterial RNA. In TRIF^+^/^+^ cells, infection with Cb NMII suppressed both NF-κB and IRF-I activation relative to DotA, consistent with earlier observations (Fig 3A and 3B). In contrast, TRIF^−^/^−^ cells showed markedly reduced responses under all tested conditions, with at least a 2-fold decrease compared to parental controls. Notably, Cb RNA treatment of TRIF^+^/^+^ cells induced robust NF-κB (1.5-3-fold) and IRF (10-12-fold) activation, whereas TRIF^−^/^−^ cells failed to respond to any tested conditions (Fig 3C and D), confirming that Cb RNA signaling in this system is TRIF-dependent. We next investigated whether Cb LPS contributes to TRIF-dependent signaling and whether suppression of TLR3/TRIF signaling is conserved in the virulent NMI strain. THP-1 Dual™ hTLR3 cells were infected with Cb NMI, NMII, or DotA strain, with purified Cb NMI LPS tested in parallel. Purified Cb NMI LPS did not induce NF-κB activation, whereas both DotA infection and *E. coli* LPS elicited strong signaling responses (Fig 4). This demonstrated Cb LPS does not contribute to host NF-κB activation in this system and that PRR signaling modulation is mainly driven by T4SS-dependent mechanisms. Accordingly, activation of TRIF-dependent pathways downstream of TLR4 do not contribute to this observed phenotype, supporting the conclusion that Cb RNA is a dominant PAMP driving TLR3/TRIF-dependent responses (Fig 3C, 3D, and 4). Collectively, these findings demonstrate that Cb employs its T4SS to modulate host TLR3/TRIF-mediated NF-κB-dependent cytokine responses and IRF-dependent IFN-I signaling. Furthermore, Cb RNA functions as a potent PAMP capable of activating these pathways, suggesting that T4SS-mediated inhibition of TLR3/TRIF signaling is a key immune evasion strategy that enables intracellular survival and replication.

**Fig 3.**
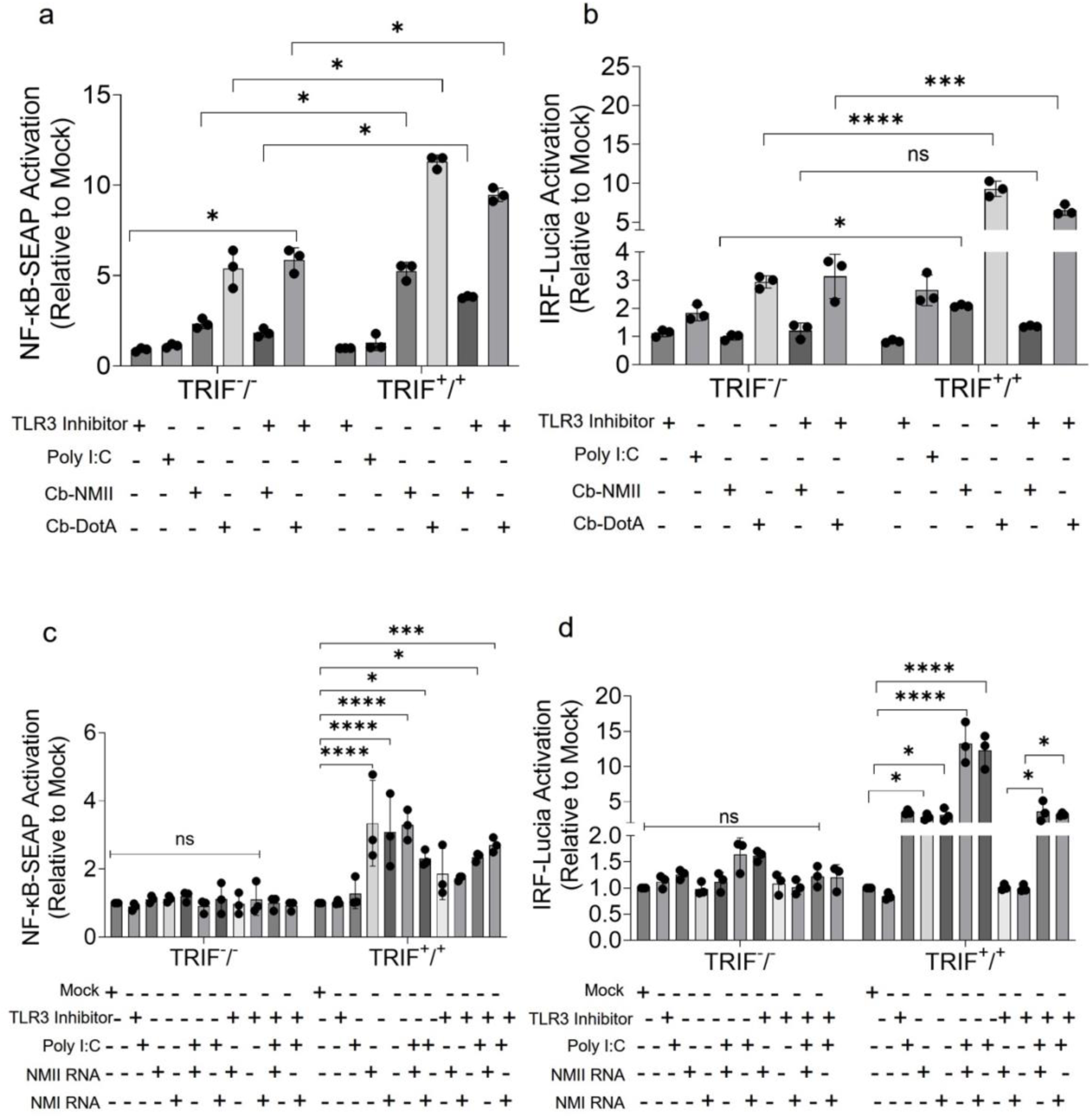
Cb suppresses NF-κB and IRF signaling via TRIF in a T4SS-dependent manner. (a-d) THP-1-Dual-TRIF-KO and THP-1-Dual (control) cells were infected with wild-type Cb NMII or the T4SS-deficient mutant DotA at an MOI of 25 for 3 h, or treated with RNA (1 µg/well) purified from virulent NMI or avirulent NMII. Cells were then stimulated with the TLR3 agonist poly(I:C) (20 μg/ mL) with or without the TLR3 inhibitor thiophene-carboxamidopropionate (10 μg/ mL). NF-κB and IRF activation were quantified in cell culture supernatants by Lucia luciferase and SEAP assays, respectively. Activation is expressed as fold change relative to mock-treated cells (set to 1). Data is expressed as mean ± SD from three independent experiments, each performed in technical pentaplicates (*n* = 5 per condition). Statistical analysis: two-way ANOVA followed by Šídák’s multiple comparisons test. Significance: *, *p* < 0.05; **, *p* < 0.01; ***, *p* < 0.001; ****, *p* < 0.0001; ns, not significant, *p* ≥ 0.05.

**Figure 4.**
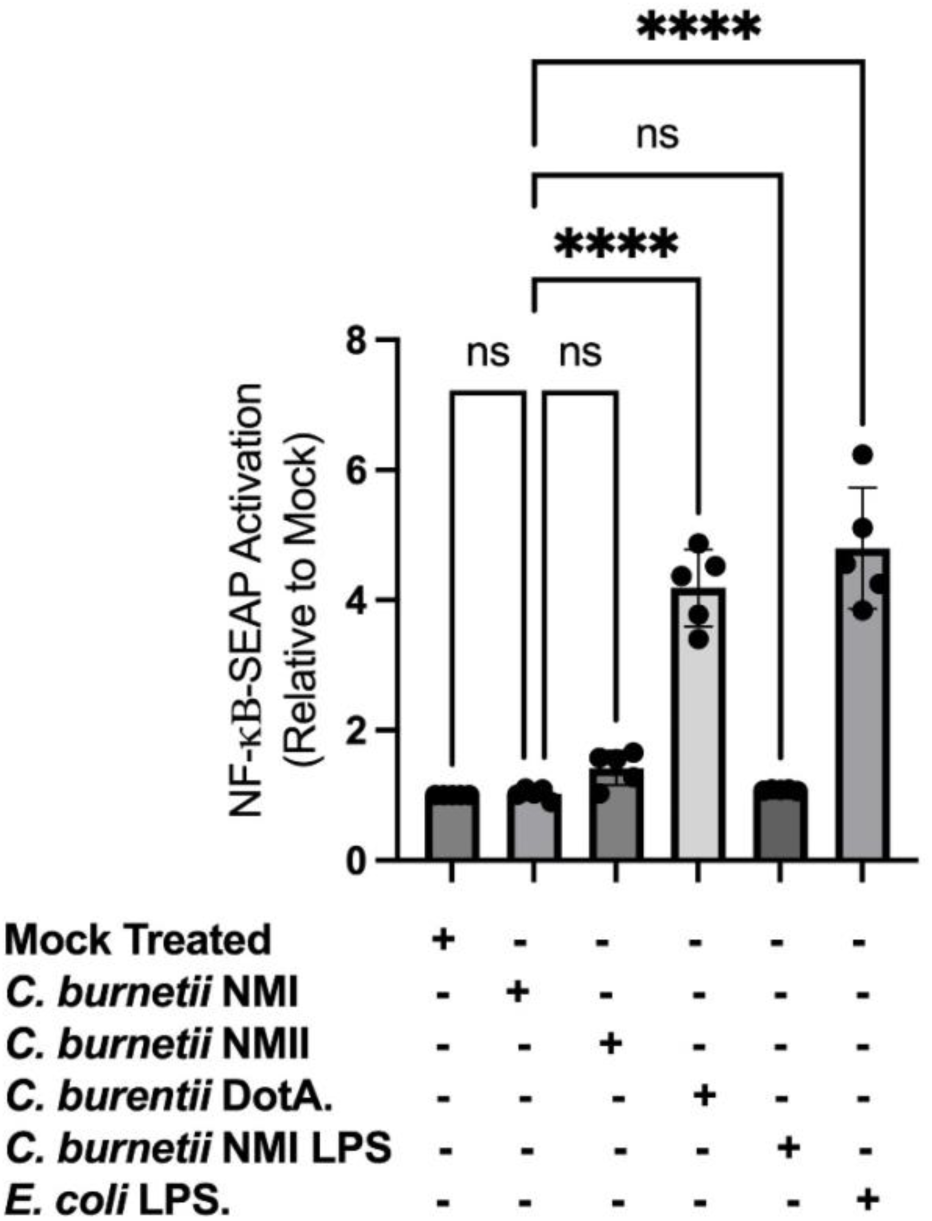
Cb inhibits host NF-κB activation through its T4SS independent of LPS stimulation. THP-1-Dual™-hTLR3 cells were either infected (MOI = 25) with virulent Cb NMI, avirulent NMII, or NMII DotA mutant, or treated with purified NMI LPS (2.5 ng/ml) or *E. coli* LPS (2.5 ng/ml; positive control). NF-κB activation was quantified using a SEAP reporter assay. Both NMI and NMII Cb infections suppressed NF-κB activation, whereas infection with the NMII DotA mutant induced robust NF-κB signaling. In contrast, purified Cb NMI LPS failed to activate NF-κB, indicating that NF-κB modulation during infection is not mediated by Cb LPS but instead depends on T4SS-driven immune suppression. Activation is expressed as fold change relative to mock-treated cells (set to 1). Data is a representative of 1 of 3 independent experiments as mean ± SD, each performed in technical pentaplicates (*n* = 5 per condition). Statistical analysis: one-way ANOVA with Dunnets multiple comparisons for all conditions against NMI.

### Both TLR3/TRIF-dependent and IFNAR-dependent immune pathways contribute to host defense against Cb

Previous studies using mice deficient in several TLRs and IFNAR have demonstrated the impact of these pathways on Cb disease severity. Both *in vitro* and *in vivo* studies have implicated TLR2 in Cb pathogenesis (16, 24, 40), while studies using IFNAR^−^/^−^ mice infected with virulent NMI suggests IFN-I signaling restricts bacterial dissemination in the peripheral tissues but not pulmonary infection (18, 41). To directly examine the contribution of the TLR3/TRIF axis *in vivo*, and distinguish it from IFNAR-dependent immunity, we sought to separate the specific contributions of TLR3-mediated signaling from downstream type I interferon responses. We employed an intra-tracheal pulmonary infection model using wild-type C57BL/6 mice (BL6-WT) and mice deficient in TLR3 (B6;129S1-*Tlr3^tm1Flv^*/J; TRL3^-/-^), TRIF (C57BL/6J-*Ticam1^Lps2^*/J; TRIF^-/-^), or IFNAR (B6(Cg)-*Ifnar1^tm1.2Ees^*/J; IFNAR^-/-^) on the same genetic background. We hypothesized that TRIF deficiency might result in a more severe phenotype than TLR3 deficiency, as TRIF also mediates signaling downstream of endosomal TLR4 and may contribute additional protection through this pathway. Mice were infected intratracheally with 1 x 10^7^ genome equivalents (GE) of virulent NMI and monitored every 2 days for disease progression by measuring body weight loss. As expected, BL6-WT mice tolerated this inoculum without losing more than 20% body weight over the course of infection (Fig 5A). In contrast, the majority of TRL3^-/-^ mice exhibited rapid and severe weight loss, reaching ≥20% loss by 7day post-infection (dpi) and therefore requiring euthanasia according to AUP guidelines. Consequently, the study was terminated on day 7 rather than the planned 14-day endpoint (Fig 5A). Several TRIF^-/-^ mice exhibited ≥20% body weight loss, disease severity within this group was more variable (Fig 5A). These findings indicate that TLR3 signaling plays a critical role in controlling disease severity during pulmonary NMI infection and that loss of this pathway results in pronounced cachexia. Furthermore the addition of poly:IC to Cb infected THP-1 cells in vitro was detrimental based on vacuole size suggesting that uptake of poly:IC and activation of TLR3 is inhibitory to Cb at a cellular level (S9 Fig). Consistent with these observations, both TRL3^-/-^ and TRIF^-/-^ mice exhibited higher pulmonary bacterial burdens compared to BL6-WT and IFNAR^-/-^ mice (Fig 5C). We speculate that lung bacterial burdens and histopathological severity in the TRL3^-/-^ and TRIF^-/-^ groups would likely have been substantially greater than the BL6-WT control had the experiment proceeded to the 14-day course. Interestingly, IFNAR^-/-^ mice displayed significantly greater splenomegaly, despite exhibiting less severe pulmonary inflammation than BL6-WT mice (Fig 5B and 5D). H&E-stained lung sections also revealed variable degrees of moderate-to-severe lymphohistiocytic interstitial inflammation across all infected groups, frequently coalescing into multifocal regions of pulmonary consolidation (Fig 5D and 5E). Collectively, these results demonstrate that TLR3/TRIF-dependent immunity plays a crucial role in restricting bacterial replication within the lung and limiting disease severity during pulmonary Cb infection, whereas IFNAR-dependent signaling appears to contribute more prominently to restricting systemic dissemination.

**Figure 5.**
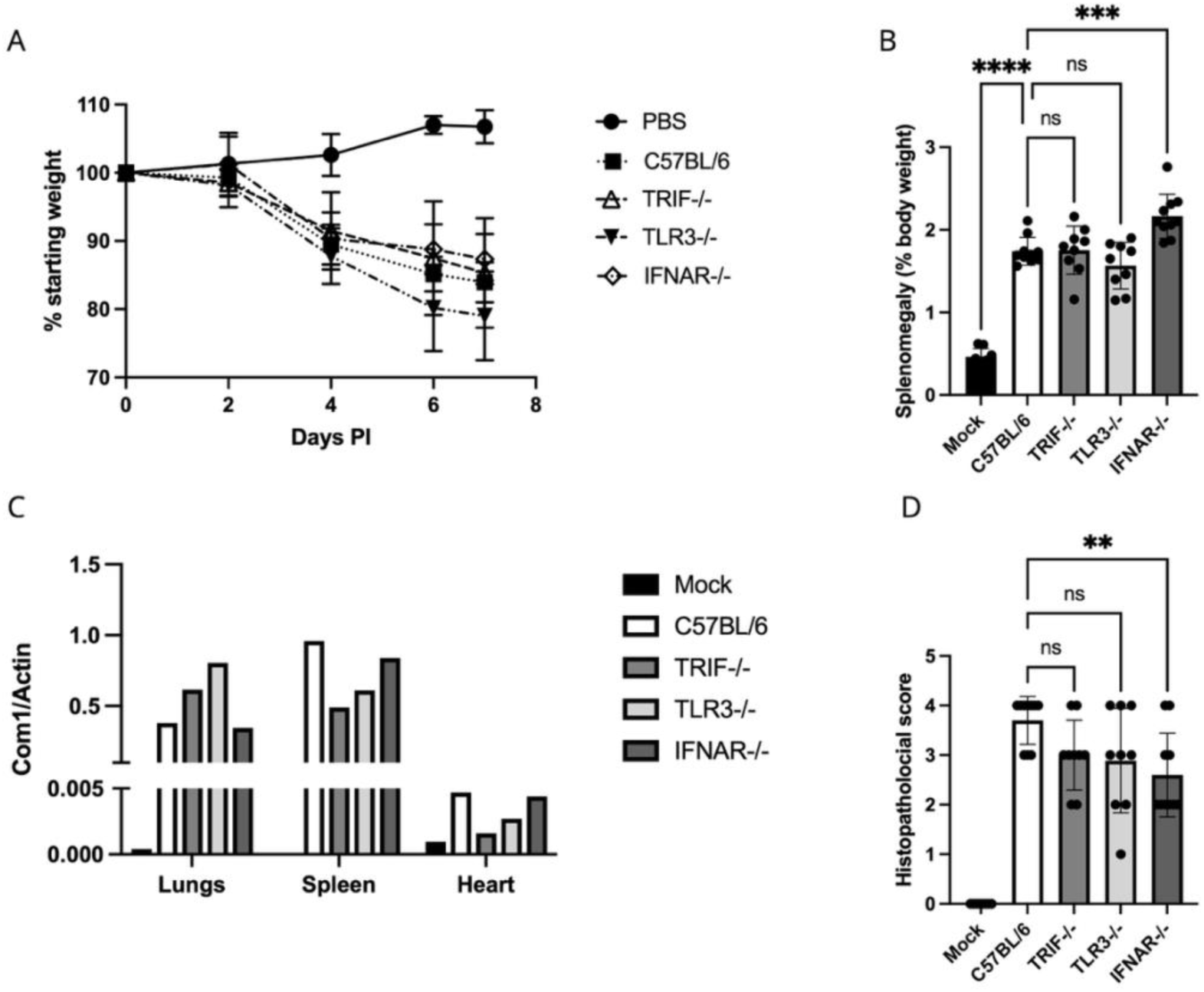
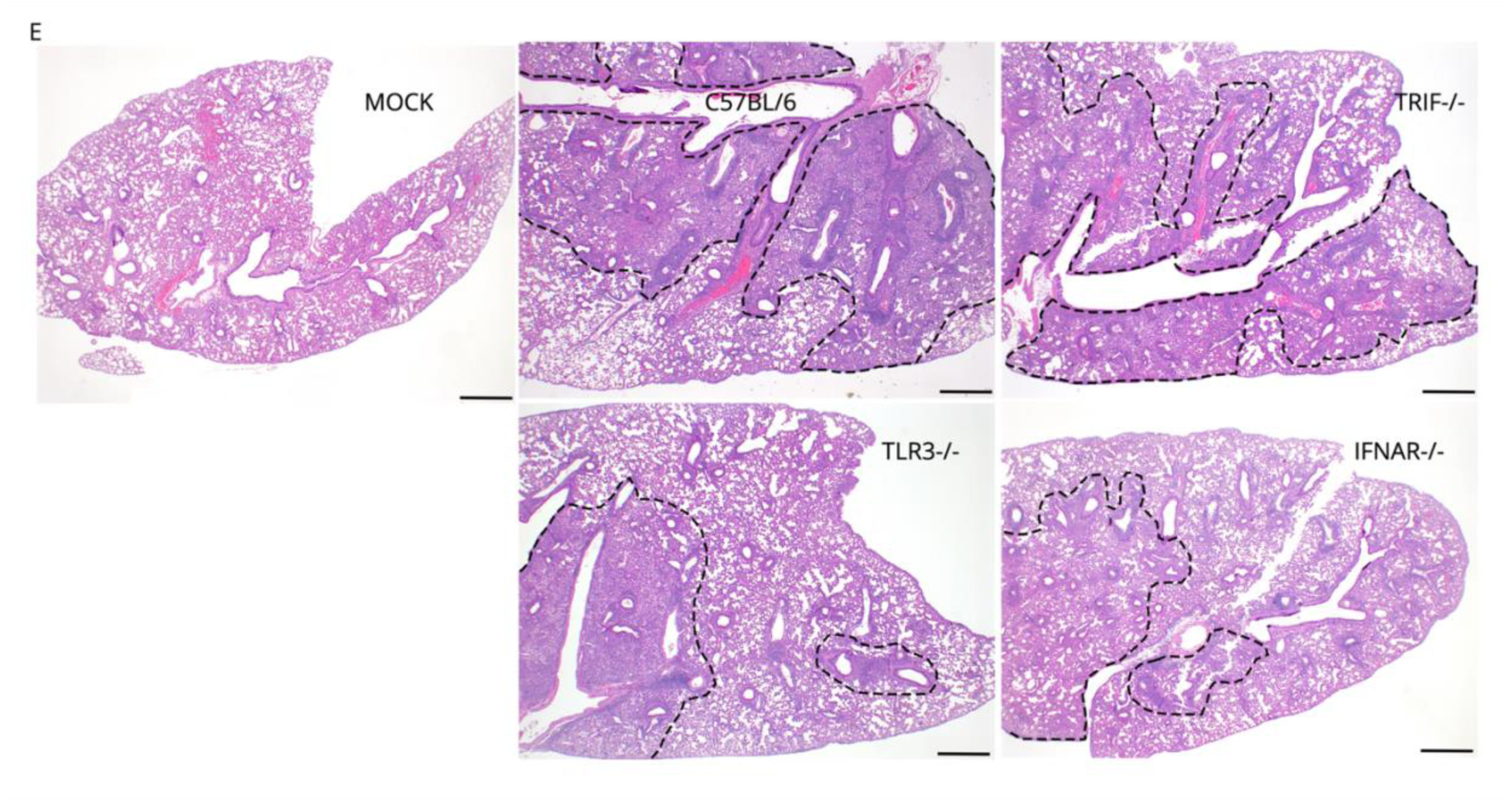
Cb NMI infection in C57BL/6 and innate immune signaling pathway deficient mice. C57BL/6-WT (strain 000644), TLR3^−/−^ (strain 005217), TRIF^−/−^ (strain 005037), and IFNAR^−/−^ (strain 028288) mice (Jackson Laboratory) were infected via the intra-tracheal route with 1 x 10^7^ GE of Cb NMI. (A) Body weight was recorded every 2 days and expressed as a percentage of the starting weight (two independent experiments, n = 5 mice/group per experiment). (B) splenomegaly assessed at necropsy (at 7 dpi) and expressed as spleen weight relative to total body weight. (C) Bacterial burden in pooled lung, spleen, and heart tissues was quantified by digital droplet PCR and expressed as NMI genome copies normalized to mouse β-actin. (D) Lung histopathology scores for individual mice. (E) Representative micrographs of lung sections from each group showing moderate to severe lymphohistiocytic interstitial inflammation with multifocal areas of consolidation (delineated by dotted lines). Images were scored for severity of inflammation. H&E stained; magnification 2×; scale bar, 500 μm. Data represents mean ± SD of 10 mice (2 biological replicates of n=5 mice/group with equal numbers of male and femal mice). *p*-values were measured using ordinary one-way ANOVA with Šídák’s multiple comparisons test.

### Cb disrupts TLR3 association with CCVs in a T4SS-dependent manner during mid-infection

To elucidate the mechanisms by which Cb manipulates TLR3-mediated signaling during infection, we performed laser scanning confocal microscopy (LSCM) with Z-Stack acquisition to examine the sub-cellular localization dynamics of host TLR3. In resting cells, TLR3 predominantly localizes to cytoplasmic endosomes, consistent with previous reports (S10 Fig) (42). We analyzed TLR3 trafficking at two critical infection time points: 6 hpi, representing the earliest stage at which T4SS-dependent inhibition of TLR3 signaling was observed (Fig 1A), and 48 hpi, corresponding to late infection when CCV expansion is pronounced and extensive host cell manipulation are well-established. At 6 hpi, host TLR3 transiently associated with Cb, exhibiting colocalization with bacteria within tight, individual unfused CCVs. In contrast, by 48 hpi, TLR3 no longer colocalized with mature, replicative-CCV (Fig 6A-6C). To determine whether this phenotype was dependent on the Cb T4SS, HeLa cells were infected with the DotA mutant at an MOI of 25 and and TLR3 localization was similarly assessed at 6 and 48 hpi. In contrast to Cb NMII infection, Z-stack LSCM analysis revealed that host TLR3 remained associated with DotA-containing vacuoles at both 6 and 48 hpi (Fig 7A-7C). Collectively, these findings demonstrate that Cb employs T4SS to prevent sustained recruitment of host TLR3 to the CCV during later stages of infection, thereby limiting the ability of TLR3 to engage Cb RNA within the CCV and initiate downstream signaling responses.

**Fig 6.**
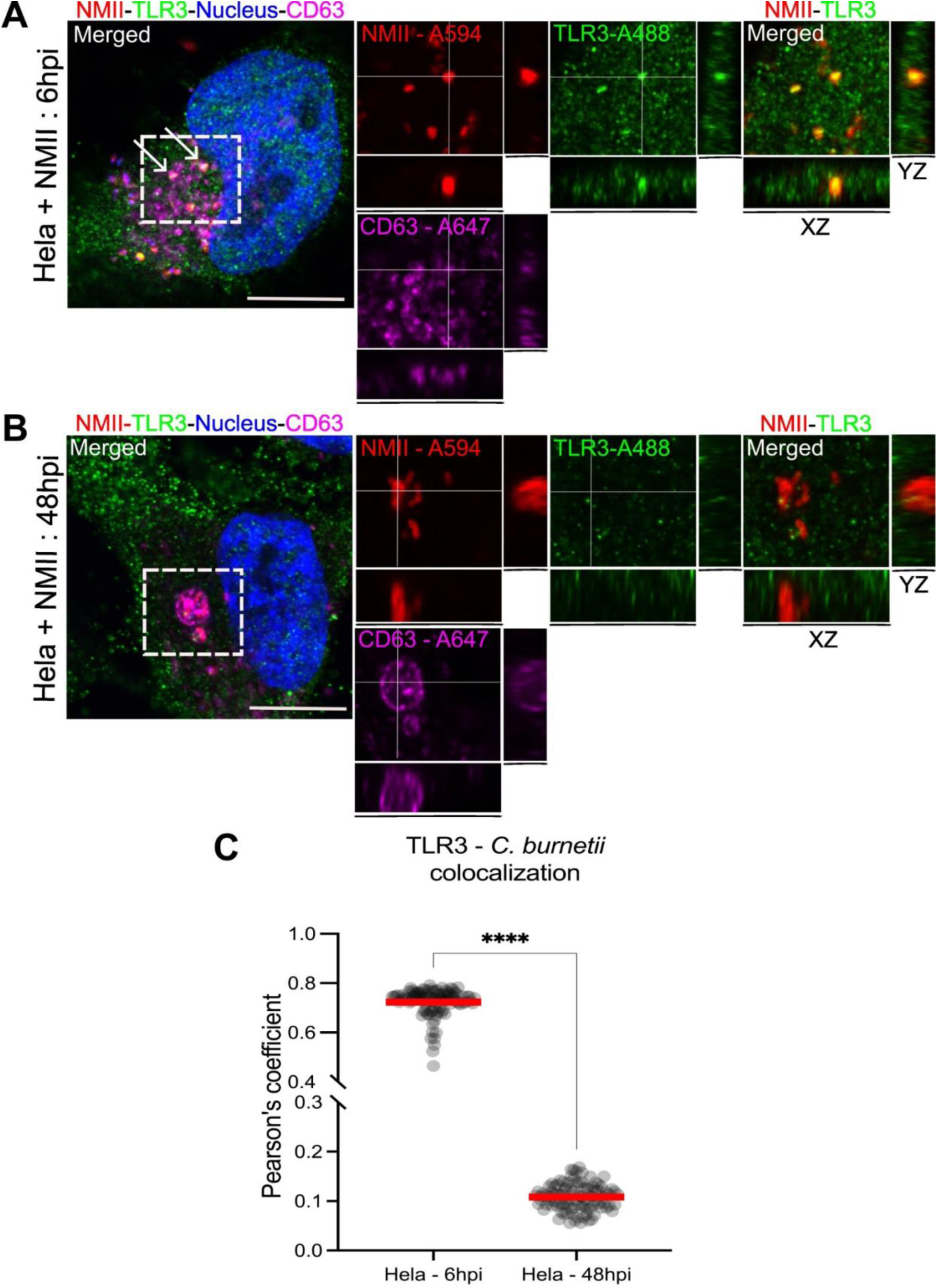
Cb effectively blocks trafficking of host TLR3 to the CCV. (A-C) LSCM Z-stack images depict the interaction between host TLR3 and wild-type NMII (MOI = 25) in a HeLa cell infection model at 6 and 48 hpi. The scale bar represents 10 μm, and colocalization is indicated by arrows. Pearson’s correlation coefficient was calculated to quantify colocalization between NMII and TLR3 (red-green overlap). (A) At 6 hpi, host TLR3 shows significant colocalization with NMII, localized within tight, unfused individual CCVs, as reflected by a high Pearson’s coefficient. (B) By 48 hpi, TLR3 no longer colocalizes with NMII, which resides in a single large, mature CCV, suggesting that Cb prevents TLR3 trafficking to the CCV during later stages of infection. (C) Quantitative colocalization analysis from three independent experiments demonstrates significant differences in TLR3-NMII colocalization over time, measured as Pearson’s coefficient for individual cells. Mean values are displayed in red, and statistical significance (p < 0.05) between colocalization at 6 hpi and 48 hpi was determined using an unpaired t-test with Welch’s correction. These findings indicate temporal regulation of TLR3 interaction with Cb.

**Fig 7.**
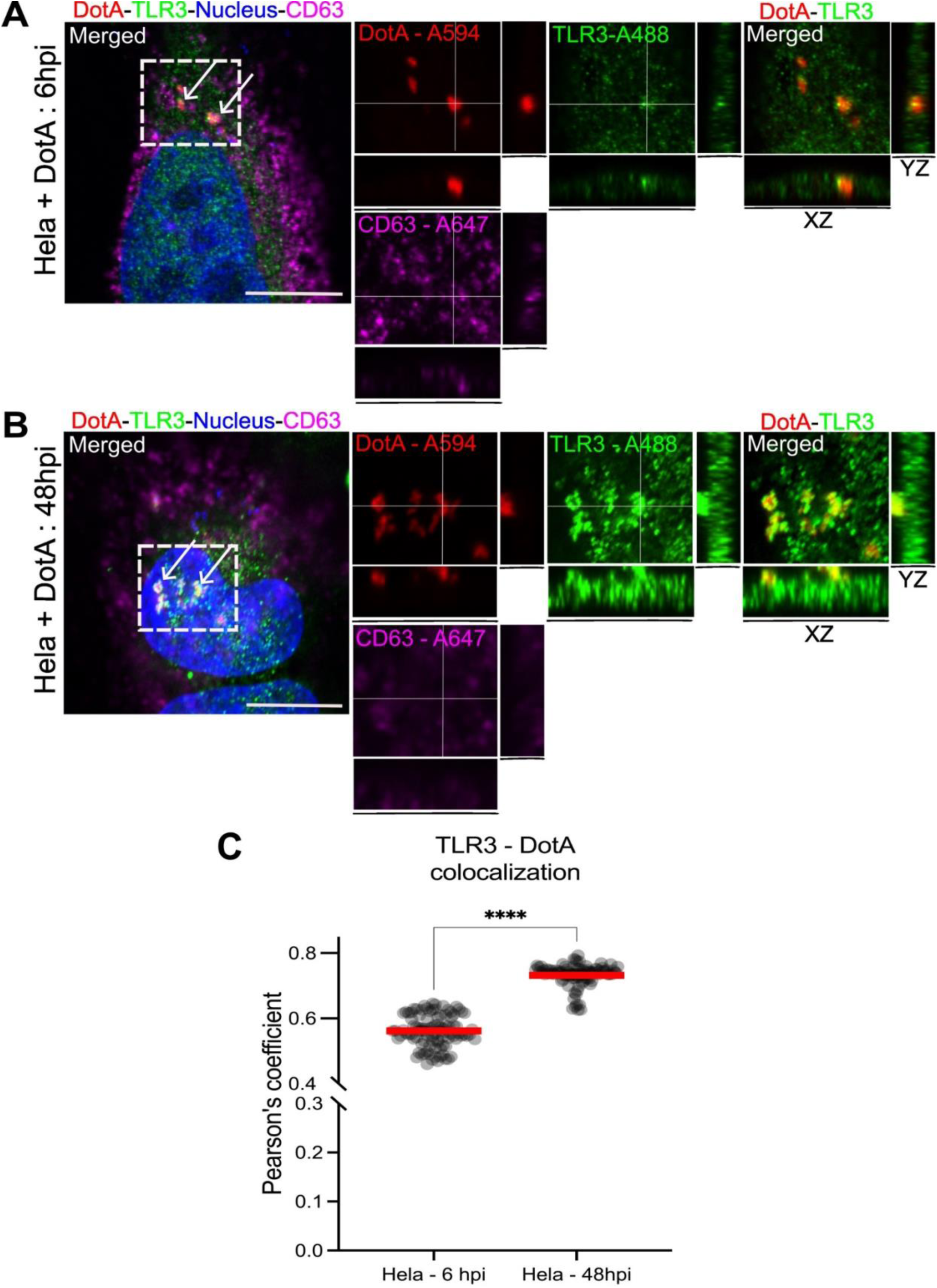
Host TLR3 co-localizes with Cb DotA during infection. (A-C) LSCM Z-stack images demonstrate the interaction between host TLR3 and DotA mutant (MOI = 25) during infection. The scale bar represents 10 μm. Colocalization of TLR3 (stained green) with DotA-containing CCVs (stained red) is evident, with areas of overlap marked by arrows, indicating that TLR3 traffics to and interacts with CCVs in the absence of a functional T4SS. (A) At 6 hpi, host TLR3 colocalizes with DotA within tight, unfused individual CCVs. (B) At 48 hpi, host TLR3 continues to colocalize with DotA-containing CCVs, in contrast to wild-type NMII, which prevents TLR3 interaction with CCVs at later stages of infection (see Fig 3). (C) A quantitative colocalization analysis of 100 cells from three independent experiments, expressed as Pearson’s coefficient, reveals sustained TLR3-DotA interaction across time points, with mean values shown in red. Statistical significance (p < 0.05) for TLR3-DotA colocalization at 6 hpi vs. 48 hpi was determined by unpaired t-test with Welch’s correction. The sustained localization of TLR3 with DotA-containing CCVs at both early and mid-stages of infection suggests that host cells attempt to recognize the pathogen through TLR3-mediated detection mechanisms, a process disrupted by the functional T4SS in Cb.

### Cb suppresses NF-κB signaling through inhibition of TRIF/TRAF6 interaction

The TLR3 signaling cascade is initiated upon receptor activation, which facilitates transient recruitment of the adaptor protein TRIF. Activated TRIF subsequently dissociates from TLR3 and engages TNF receptor-associated factor 6 (TRAF6) and receptor-interacting serine/threonine-protein kinase 1 (RIPK1) to initiate downstream NF-κB activation and induction of pro-inflammatory cytokines (32, 43, 44). Having established that Cb manipulates TLR3/TRIF signaling in a T4SS-dependent manner with Cb-RNA functioning as the PAMP, we next sought to mechanistically determine how Cb disrupts TLR3/TRIF-mediated downstream signaling responses. To examine whether Cb alters TRIF-TRAF6 engagement, we assessed the interaction between endogenous TRIF and ectopically expressed FLAG-tagged human TRAF6 in HeLa cells infected with either Cb NMII or the DotA strain. Co-immunoprecipitation (IP) assays were performed using anti-FLAG M2 affinity agarose to capture FLAG-TRAF6 and associated host proteins.

IP analysis showed that wild-type Cb NMII infection inhibited the interaction between endogenous TRIF and FLAG-TRAF6, while DotA-infected cells exhibited increased levels of TRIF co-immunoprecipitating with FLAG-TRAF6 (Fig. 8A). Whole-cell lysates (WCLs) were also analyzed by western blotting to confirm expression of FLAG-TRAF6 and endogenous TRIF (Fig. 8B). To determine whether TLR3 pathway activation could overcome this Cb-mediated inhibition of TRIF/TRAF6, we treated NMII-infected HeLa cells with the TLR3 agonist poly(I:C). Notably, IP analysis demonstrated that Cb NMII continued to suppress the TRIF-TRAF6 interaction even following poly(I:C) stimulation (Fig. 8C). In contrast, poly I:C treatment of DotA-infected HeLa cells promoted a robust interaction between endogenous TRIF and FLAG-TRAF6, consistent with preserved TLR3/TRIF signaling in the absence of T4SS activity (Fig. 8C). Western blot analysis of WCLs confirmed equivalent expression of FLAG-TRAF6 and endogenous TRIF across all experimental conditions (Fig 8D). These findings demonstrate that Cb suppresses downstream TRIF signaling by disrupting TRIF–TRAF6 complex, thereby limiting NF-κB activation during infection.

**Fig 8.**
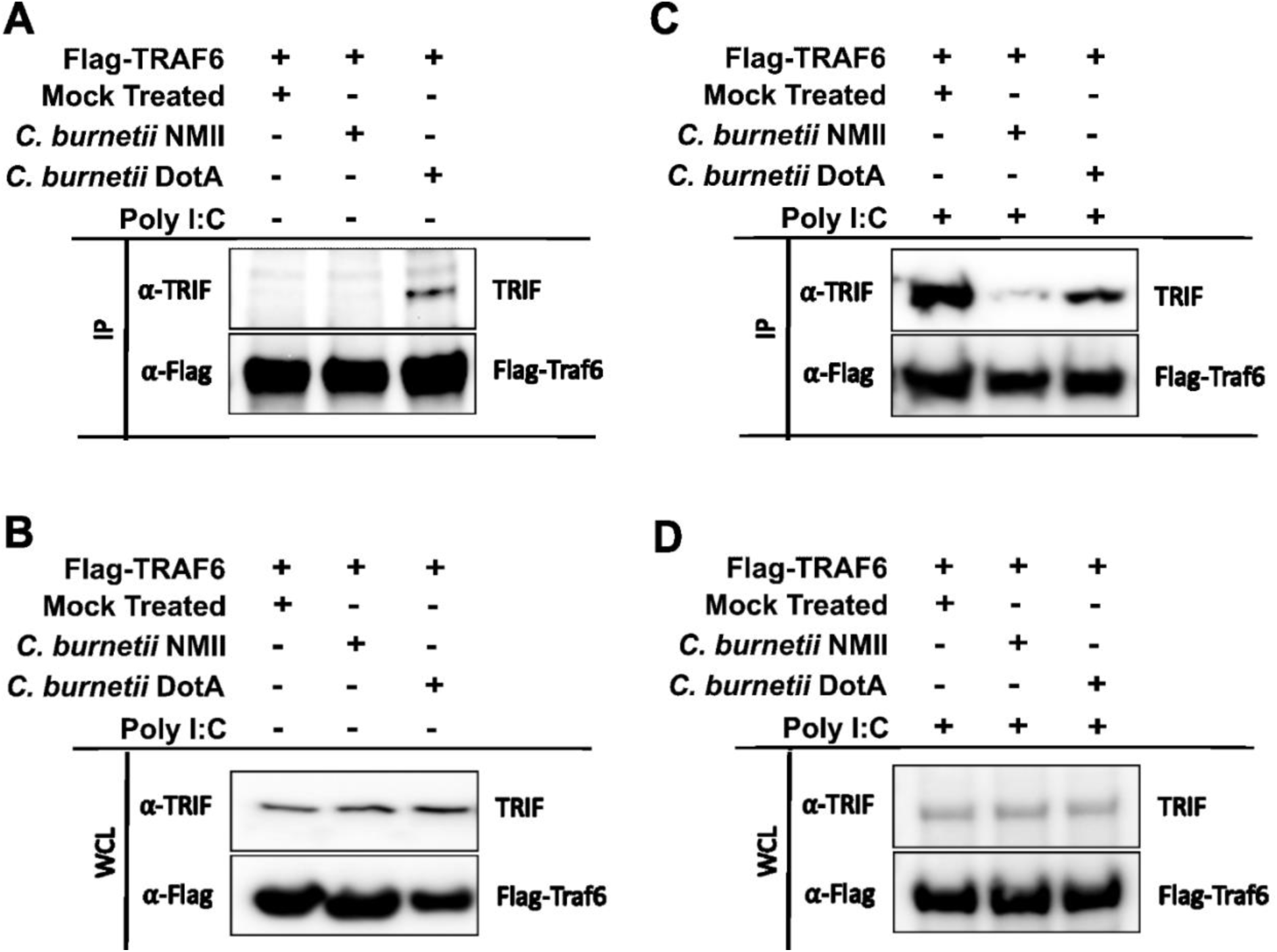
Cb T4SS-mediated inhibition of host TRIF and Flag-tagged TRAF6 interaction promotes suppression of downstream NF-κB activity. (A) Western blots showing IPs of ectopically expressed Flag-tagged human TRAF6 in HeLa cells infected with either wild-type NMII or DotA mutant. Mock-treated HeLa cells expressing Flag-tagged TRAF6 served as controls. To investigate the TRIF-TRAF6 interaction, Flag-IPs were probed for the presence of endogenous TRIF and Flag-TRAF6. The findings indicate that endogenous TRIF and Flag-TRAF6 interact robustly in HeLa cells infected with the DotA mutant, whereas this interaction is notably inhibited in cells infected with wild-type NMII. (B) Western blot analysis of whole-cell lysates (WCLs) probed with antibodies against endogenous TRIF and Flag-TRAF6 (Table S1). Expression of endogenous TRIF and Flag-TRAF6 was verified by western blotting 10% of the total input from each sample. (C) Western blots illustrating IPs of ectopically expressed Flag-tagged human TRAF6 in HeLa cells infected with either wild-type NMII or DotA mutant and stimulated with poly I:C (2 μg/ml). Mock-treated HeLa cells expressing Flag-TRAF6 and stimulated with poly I:C were used as controls. To evaluate the TRIF-TRAF6 interaction, Flag IPs were probed with antibodies against endogenous TRIF and FLag-TRAF6. The findings demonstrate that TLR3 stimulation fails to promote the interaction between endogenous TRIF and Flag-TRAF6 in HeLa cells infected with wild-type NMII. In contrast, robust interaction between endogenous TRIF and Flag-TRAF6 persists in cells infected with the DotA mutant. (D) HeLa cells ectopically expressing Flag-tagged human TRAF6 were mock-treated or infected with wild-type NMII or the DotA mutant, followed by stimulation with poly I:C (2 μg/ml). WCLs were analyzed by Western blot for endogenous TRIF and Flag-TRAF6 (Table S1), and 10% of each lysate was reserved as an input control to verify protein expression prior to Flag-IP.

### Multiple Cb T4SS effectors suppress host NF-κB signaling, with CBU1292 contributing to modulation of both NF-κB and IRF activation

To identify Cb T4SS effectors involved in inhibition of NF-κB signaling, a Himar1 transposon mutant library containing single effector insertion mutants was screened for the ability to induce NF-κB luciferase activity in THP-1 Lucia reporter cells relative to the Cb NMII strain. Among the 32 effector mutants screened, five candidates significantly induced NF-κB activation suggesting that Cb could dampen NF-κB signaling through multiple effectors including CBU0021 (*cvpB/cig2*), CBU1699, CBU1292 (*ankK*), CBU0201 (*ankC*), and CBU1254(S10 Fig) (one-way ANOVA with Dunnett’s test, *p* < 0.05). A representative subset of the screen, including these five T4SS effector candidates and the previously characterized NF-κB-modulating effector CBU1639 (20) is shown in Fig 9A. Interestingly, among the five mutants identified, CBU1292::Tn (*ankK::Tn*) also induced IRF reporter activation, suggesting a role in regulation of IFN-I signaling. Complementation of the CBU1292::Tn mutant (S11A and S11B Fig) restored inhibition of both NF-κB and IRF activation to levels comparable to those observed in Cb NMII-infected cells (Fig. 9B and 9C). Mechanistically, the mutant showed ability to block poly I:C induced activation of reporter activity, indicating that CBU1292 targets signaling downstream of TLR3 stimulation. Collectively, these findings highlight a critical role for the T4SS in coordinated regulation of NF-κB and IRF-dependent signaling and identify CBU1292 (ankK) as a dual regulator of both pathways downstream of the TLR3 axis.

**Fig 9.**
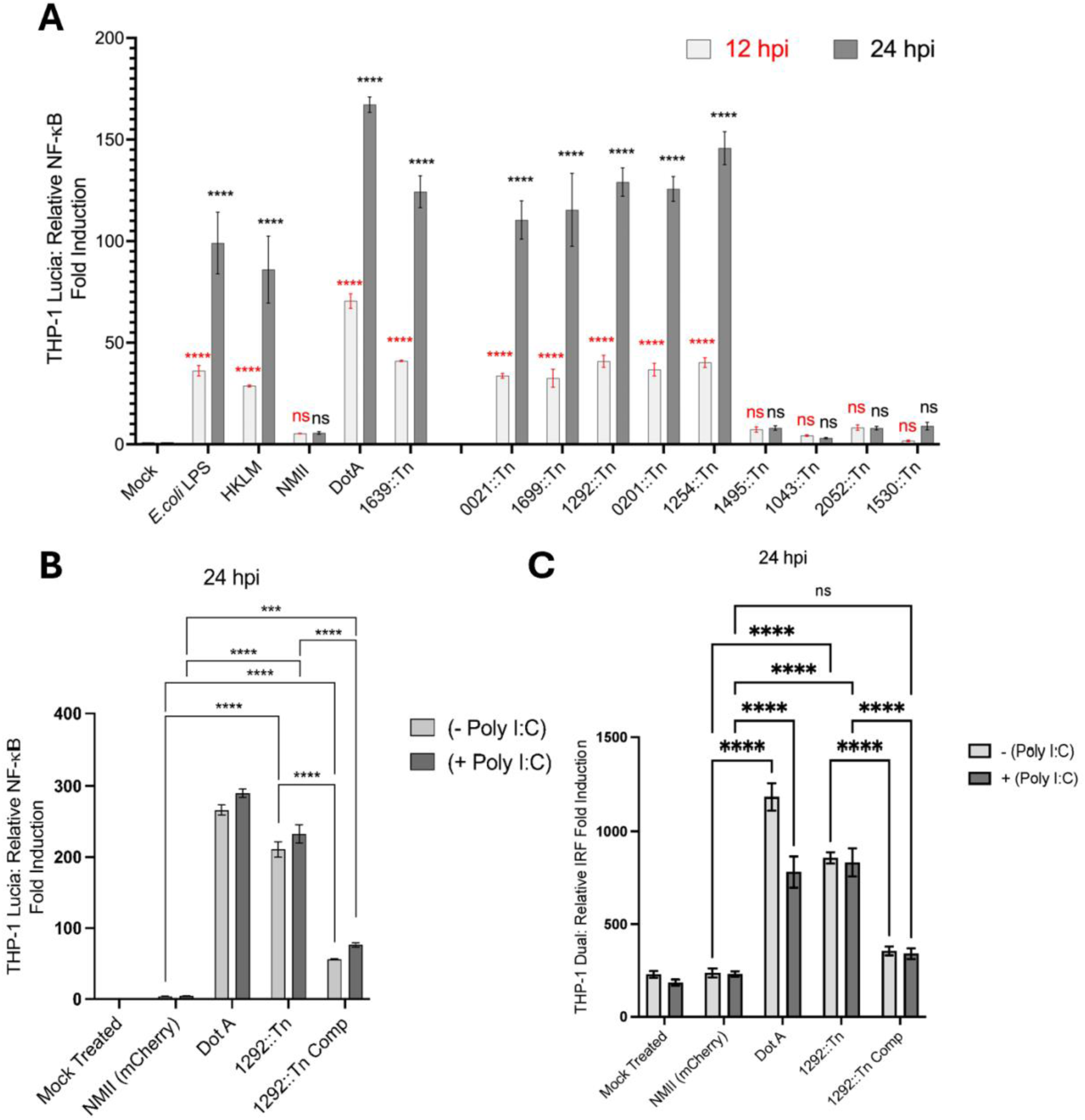
Multiple T4SS effectors contribute to the disruption of host TLR3/TRIF-induced NF-κB activation by Cb. (A) A subset of five Cb T4SS effector candidates were identified as inhibitors of NF-κB activation in THP-1 Lucia cells. Compared to mock treated cells, relative luciferase activation levels were significantly elevated in the DotA mutant, T4SS effector positive control CBU1639::Tn, as well as in the candidate strains CB*U1254*::Tn, CB*U1292*::Tn (*ank*K::Tn), CB*U0201*::Tn (*ank*C::Tn), CB*U0021*::Tn (*cvp*B::Tn or *cig*2::Tn), and CB*U1699*::Tn, at both 12 hpi and 24 hpi. NF-κB-Lucia activation levels in wild-type NMII-infected cells were similar to mock-treated cells. Data are presented as mean ± SEM from three independent experiments. Statistical comparisons were performed using one-way ANOVA with Dunnett’s test for multiple comparisons between mock-treated THP-1 Lucia cells and those treated with *E. coli* LPS or HKLM, or infected with wild-type NMII, or the DotA mutant, or NMII T4SS effector candidate Tn mutant strains. Error bars represent SEM, with statistical significance (p < 0.05) at 12 hpi highlighted in red and at 24 hpi indicated in black. (B) Complementation of CB*U1292*::Tn (*ankK*::Tn) strain suppresses NF-κB activation at 24 hpi. Compared to cells infected with mCherry-expressing wild-type NMII, NF-κB-Lucia activation levels were higher in cells infected with CB*U1292*::Tn (*ank*K::Tn). However, inoculation of the complemented strain CB*U1292*::Tn comp (*ank*K::Tn comp) into THP-1 NF-κB-Lucia cells resulted in NF-κB-Lucia inhibition. The addition of poly I:C (20µg/ml) to cells infected with the complemented strain did not rescue NF-κB-Lucia inhibition. The data represents the mean ± SEM of three independent experiments. Error bars and p-values < 0.05 are highlighted for comparisons. Statistical significance was determined using two-way ANOVA with Tukey’s test for multiple comparisons. (C) Complementation of *Cbu1292*::*Tn* (*ankK*::*Tn*) strain suppresses IRF activation at 24 hpi. Compared to cells infected with mCherry-expressing wild-type NMII, IRF-Lucia activation levels were higher in cells infected with CB*U1292*::Tn (*ank*K::Tn). However, inoculation of the complemented strain CB*U1292*::Tn comp (*ankK::Tn* comp) into THP-1 IRF-Lucia cells resulted in IRF-Lucia inhibition. The addition of poly I:C (20µg/ml) to cells infected with the complemented strain did not rescue IRF-Lucia inhibition. The data represents the mean ± SEM of three independent experiments. Error bars and p-values < 0.05 are highlighted for comparisons. Statistical significance was determined using two-way ANOVA with Tukey’s test for multiple comparisons.

## Discussion

Many host defense pathways are activated by host sensors that detect microbial determinants introduced into host cells during infection, including those delivered by bacterial secretion systems (45). Accordingly, successful intracellular pathogens have evolved multi-faceted mechanisms to counteract these defenses and promote their survival. Following inhalation, Cb is phagocytosed by macrophages, where it establishes a replicative niche within the CCV through its T4SS (1). Notably, *in vitro* infection proceeds with minimal activation of innate immune signaling pathways, despite the presence of numerous Cb-associated PAMPs (13). This observation suggests that Cb actively suppresses host innate immune responses. Such suppression is likely essential to balance inflammatory responses or prevent host cell death, which is important to CCV expansion (46).

One mechanism contributing to immune evasion may involve sequestration of bacterial PAMPs within the CCV, limiting exposure of plasma membrane and cytosolic PRRs to immunostimulatory molecules such as LPS and peptidoglycan. Additionally, emerging evidence indicates that bacterial nucleic acids can also act as potent PAMPs, activating endosome-associated receptors including TLR3, TLR7, and TLR9, thereby shaping host cytokine and interferon responses (45, 47, 48). In this study, we show *Cb* employs its T4SS to modulate host TLR3 signalling and reveal that *Cb* specifically suppresses TLR3/TRIF-induced NF-κB and IRF-I signaling in a T4SS-dependent manner. Furthermore, we identified purified RNA from both avirulent NMII and virulent NMI Cb strains as a potent activator of TLR3/TRIF-dependent signaling, establishing Cb RNA as a biologically relevant PAMP capable of engaging this innate immune pathway. Depending on the pathogen, nucleic acid sensing has been previously reported to either promote protective host immunity or contribute to immunopathology (49). For example, *Listeria monocytogenes* secretes the RNA-binding protein Zea to deliver bacterial RNA to the cytosolic RNA sensor RIG-I, enhancing IFN-I production (50). Similarly, *Brucella abortus* nucleic acids, including RNA, induce TLR3- and TLR7-dependent IFN-I responses, although these receptors appear dispensable for *in vivo* infection control (51). In contrast, *Borrelia burgdorferi* RNA activates IFN-α and IFN-λ1 via a TLR7–IRF7 pathway (52). *Legionella pneumophila* activates TLR3, triggering downstream NF-κB signaling and induction of IL-6 and TNFα (53). Together, these findings highlight that bacterial RNA sensing is a biologically relevant pathway that may influence infection outcomes in a pathogen-specific manner.

Interestingly, we observed basal NF-κB and IRF signaling in the presence of a TLR3 inhibitor. Considering the extensive crosstalk between innate immune signaling pathways, this residual activation may reflect engagement of other PRRs, such as TLR2-dependent signaling, which is TRIF-independent and has previously been implicated in Cb innate immune recognition. Additionally, we show that purified NMI LPS failed to induce NF-κB activation in reporter cells, indicating that Cb LPS does not trigger TLR4-dependent signaling and is unlikely to account for the observed background activation. Collectively, these findings support the conclusion that Cb-driven modulation of TRIF signaling occurs predominantly through TLR3 rather than TLR4.

Consistent with these *in vitro* findings, pulmonary infection of C57BL/6 mice and knockout strains deficient in TLR3, TRIF, or IFNAR revealed distinct contributions of these pathways to host defense. TLR3^-^/^-^ mice exhibited severe cachexia and reached humane endpoint criteria during infection, necessitating early termination of the experiment. TRIF^-^/^-^ mice also demonstrated increased susceptibility, although with greater variability. In contrast, IFNAR^-^/^-^ mice displayed reduced pulmonary inflammation but increased splenomegaly, suggesting enhanced dissemination. These results demonstrate that TLR3/TRIF-dependent signaling is critical for controlling disease severity and bacterial burden in the lung, whereas IFNAR-dependent signaling plays a dominant role in restricting systemic dissemination. Together, these *in vivo* data demonstrate that TLR3/TRIF- and IFNAR-dependent immune responses, although with distinct pathological outcomes, ultimately converge to promote host protection during Cb infection and thus represent key targets for pathogen-driven immune evasion.

Importantly, our *in vivo* results reinforce the conclusion that LPS-driven TRIF signaling does not play a dominant protective role during pulmonary infection. Specifically, TLR3^-^/^-^ mice exhibited more severe disease than TRIF^-^/^-^ mice. If endosomal TLR4–TRIF signaling were a major contributor to infection control, TRIF^-^/^-^ mice would be expected to display a more severe phenotype than TLR3^-^/^-^ animals. Instead, these results are consistent with a model in which TLR3 is the primary driver of TRIF-mediated protection during pulmonary infection with virulent Cb.

Our results indicate that suppression of TLR3/TRIF signaling is crucial for Cb survival and replication *in vitro*, as TLR3-specific activation significantly impairs Cb growth in macrophages. Mechanistically, we propose that Cb inhibits TLR3 signaling through various temporally distinct, T4SS-dependent strategies. Early during infection, effectors such as CBU1292 may play a crucial role to suppress both NF-κB and IRF activation, potentially by targeting TRIF or an upstream signaling node. Cb disrupts NF-κB signaling by inhibiting the TRIF–TRAF6 interaction in a T4SS-dependent manner, whether this is a specific function of CBU1292 or another effector remains to be determined. At later time points (by 48 hpi), Cb prevents association of host TLR3 with the mature CCV in a T4SS-dependent manner. Collectively, the presence of temporally distinct mechanisms targeting the same innate immune pathway strongly suggests that suppression of TLR3/TRIF signaling is critical for Cb intracellular survival and disease progression. However, further work is needed to identify the specific Cb T4SS effector(s) responsible for these distinct modes of TLR3 pathway modulation.

In addition to CBU1292, we identified four other candidate T4SS effectors capable of modulating NF-κB signaling, further supporting the broader concept that Cb employs a multi-effector strategy to modulate host innate immune responses. Such a strategy is consistent with immune evasion mechanisms employed by numerous bacterial pathogens, which use secretion systems (including T3SS and T4SS) to disrupt multiple signaling nodes within TLR-mediated immune cascades (54–59). The study of Cb T4SS effectors remains an emerging area, with several known effectors including CinF, NopA, and CBU1639 have already been implicated in disrupting NF-κB signaling through distinct mechanisms such as phosphatase activity (CinF), interference with nucleocytoplasmic transport (NopA), and yet unidentified mechanisms (CBU1639) (13, 26, 27). Among the newly identified effectors described here, CBU1254, CBU1292 (*ank*K), and CBU0201 (*ank*C) encode ankyrin repeat domain (ARD)-containing proteins, while CBU0021 (*cvp*B, *cig*2) and CBU1699 harbor coiled-coil domains (CCDs), suggesting mechanistic diversity in how these proteins may target host signaling pathways. Notably, ARD and CCD-containing effectors from other intracellular pathogens such as *Orientia tsutsugamushi* and *L. pneumophila* have been linked to NF-κB modulation (60, 61). Further studies are needed to clarify how these Cb T4SS effectors disrupt TLR3/TRIF-induced NF-κB and IRF signaling.

Finally, previous studies have shown that TLR3^−^/^−^ mice infected with *Chlamydia muridarum* exhibit increased bacterial burden, altered cytokine profiles, and impaired recruitment of protective CD4^+^ T cells (62). Although genital tract infection differs substantially from pulmonary infection, these observations support a broader role for TLR3-dependent immune sensing in restricting intracellular bacterial pathogens. Together with our findings, these studies suggest that TLR3-mediated recognition of microbial nucleic acids may represent an underappreciated axis of antibacterial immunity that is actively targeted by intracellular pathogens such as Cb.

## Materials and Methods

### Bacterial strains and mammalian tissue culture

Cb Nine Mile phase II RSA439, clone 4 (NMII) stocks were grown in Hela cells and purified as previously described (63). The NMII T4SS-deficient *ΔdotA* mutant strain (DotA) was generously provided by Drs. Paul Beare and Bob Heinzen (NIAID Rocky Mountain Laboratories, Montana) and grown as previously described (64). To identify Cb T4SS effectors capable of inhibiting NF-κB, 32 NMII T4SS effector transposon (Tn) mutant strains from a Himar1 Tn mutant clone library were grown in ACCM media as previously described (36). For animal intratracheal infection, Cb Nine Mile phase I RSA493 clone 7 (NMI) was grown in embryonated yolk sacs and purified using density gradient centrifugation as previously described (65). Experiments involving NMI were performed in biosafety level 3 (BSL3) facilities at the Texas A&M Health Science Center, Bryan. All eukaryotic cell lines used in this study (S1 Table) were grown following either Invivogen or ATCC guidelines. Primary bone marrow–derived macrophage (BMDM) cells were prepared from C57BL6 mice (JAX) using recombinant murine m-CSF (Peprotech) as previously described (66, 67).

### Toll-Like Receptor screening assay

To determine whether Cb*’*s functional T4SS suppresses Toll-Like Receptor (TLR)-induced NF-κB activity, human monocytic THP-1 NF-κB-Lucia reporter cells were utilized according to Invivogen’s protocols. For TLR-stimulation, and NF-κB-induced Lucia luciferase reporter expression, THP-1 Lucia cells were first infected with either wild-type NMII or the T4SS-deficient DotA mutant in a 24-well tissue culture plate at a multiplicity of infection (MOI) of 25. Bacteria were added to 1 × 10^6^ THP-1 Lucia cells per well and incubated at 37°C for 2 hours. At 2 hpi, agonists targeting MyD88-dependent TLRs (TLR2, TLR4, and TLR7/8) or TRIF-dependent TLRs (TLR3 and TLR4) were introduced to NMII- or DotA-infected cells, and NF-κB-Lucia induction levels were measured at 12 hpi. A range of concentrations of well-established TLR agonists, including PAM3CSK4 (TLR2), *E. coli* LPS (TLR4), CL075 (TLR7/8), and poly I:C (TLR3), were applied to evaluate dose-response relationships (S1 Table). For NF-κB-inducible Lucia luciferase readout, cell culture supernatants from the samples were collected at 12hpi, and luminescence activity was measured using InvivoGen’s QUANTI-Luc coelenterazine-based luminescence assay. PerkinElmer EnVision 2104 multilabel reader was used to measure luminescence and NF-κB fold induction levels were calculated from a total of three independent experiments. The NF-κB fold induction level data were normalized to the mock-treated control.

### qRT-PCR analysis of host NF-κB-inducible mRNA

1 × 10^6^ THP-1 cells were infected with NMII in a 24-well tissue culture plate at a multiplicity of infection (MOI) of 25. At 2hpi, agonists for TLR3 (poly I:C) and TLR4 (LPS, *E. coli*) were added at the indicated concentrations. At 12hpi, total RNA was extracted from the infected & treated THP-1 cells using TRIzol (Invitrogen). Harvested total RNA was treated with RNase-free DNase I (Roche) and a total of 500 ng of RNA was reverse transcribed using the iScript cDNA Synthesis Kit (Bio-rad). qRT-PCR reactions were performed in triplicates using PowerUp™ SYBR™ Green Master Mix (Applied Biosystems) on an Applied Biosystems QuantStudio 6. Cycle threshold (Ct) values for each NF-κB-inducible mRNA transcript (*IL8*, *TNFα*, and *BCL3*) were normalized to the expression of human *β-actin* (i.e., reference gene) and the fold changes from three independent experiments were determined by using the 2^-ΔΔCt^ method (68). qRT-PCR primers used for each mRNA transcript are listed in the S2 Table.

### Dual NF-κB (SEAP) and IRF (Lucia) reporter assays

To quantify NF-κB and IRF activation in response to Cb infection, THP-1-Dual™ hTLR3 reporter cells (InvivoGen) were seeded at 1 × 10⁶ cells/well in 24-well tissue culture plates and infected with wild-type NMII or the T4SS-deficient DotA mutant (MOI = 25) for 3 h. Supernatants were collected and NF-κB-inducible SEAP activation was measured using QUANTI-Blue™ (InvivoGen) at 635 nm (Absorbance), while IRF-inducible Lucia Luciferase activation was measured using QUANTI-Luc™ (InvivoGen) luminescence reagent on a BioTek Cytation 5 multimode reader (Agilent). Poly I:C (20 µg/mL) and thiophene-carboxamidopropionate (10 µg/mL) were used as TLR3 agonist and inhibitor controls, respectively. To assess NF-κB and IRF activation in response to bacterial RNA, wild-type THP-1-Dual™ and THP-1-Dual™ TRIF-KO cells were seeded at 1 × 10⁶ cells/well in 24-well tissue culture plates and treated with purified Cb RNA (1 µg/well) from either virulent NMI or avirulent NMII Cb strains. TLR3 agonist poly I:C, and TLR3 inhibitor thiophene-carboxamidopropionate were included as controls. NF-κB and IRF reporter activity was quantified from supernatants as described above.

### NF-κB-inducible (SEAP) reporter assay to assess Cb LPS stimulation

To evaluate whether Cb LPS contributes to NF-κB pathway activation, THP-1-Dual™ hTLR3 reporter cells (InvivoGen) were seeded at 1 × 10⁶ cells/well in 24-well tissue culture plates. Cells were infected with Cb NMI, NMII, or the T4SS-deficient NMII DotA mutant at a multiplicity of infection (MOI) of 25. In parallel, uninfected cells were treated with purified Cb NMI LPS (2.5 ng/mL) or *E. coli* LPS (2.5 ng/mL) as a positive control for NF-κB activation. Following infection or stimulation, cell culture supernatants were harvested and NF-κB activation was quantified using the NF-κB-inducible secreted alkaline phosphatase (SEAP) reporter assay. SEAP activity was measured using QUANTI-Blue™ reagent (InvivoGen), and absorbance was recorded at 635 nm using a BioTek Cytation 5 multimode plate reader (Agilent).

### Animal experiments

All the *in vivo* mice experiments performed in this study were approved by the Institutional Animal Care and Use Committee (IACUC) of Texas A&M University. The mouse strains used in this study are listed in the S1 Table. TLR3⁻/⁻, TRIF⁻/⁻, and IFNAR1⁻/⁻ mice strains were congenic on the BL6-WT background and compared directly with the BL6-WT controls. Six-eight weeks old BL6-WT, TLR3⁻/⁻, TRIF⁻/⁻, and IFNAR1⁻/⁻ mice strains were housed under pathogen-free conditions. Aerosol infections with virulent wild-type NMI Cb were performed as previously described (69). Briefly, 1 × 10^7^ GE of NMI Cb were delivered intratracheally into groups of 10 mice (5 females and 5 males per group). Mice were monitored daily for clinical signs of disease and weighed every 2 days. The study was designed to continue for 14 dpi or until mice reached predefined humane endpoints (>20% body weight loss), at which time they were euthanized in accordance with approved animal use protocols. For histopathological analysis, the left lung lobe was collected at necropsy and fixed in 10% neutral buffered formalin for 72 hours. Fixed tissues were processed by AML Laboratories (St. Augustine, FL, USA), embedded in paraffin, and sectioned at 5 μm thickness. Sections were evaluated by an ACVP board-certified veterinary anatomic pathologist. Pulmonary lesions were assessed using a semi-quantitative scoring system (0–6 scale) as described in S3 Table. Quantification of NMI genomic DNA (gDNA) in the lung, spleen, and heart was performed by digital droplet PCR (Fig 5C and <u>S12 Fig</u>) as described below.

### Cb enumeration assay

For animal experiments, Digital droplet PCR (ddPCR) was performed to quantify Cb genomic DNA (gDNA) in mouse tissues. For each experimental group, organ samples (lung, spleen, and heart) were collected in 1x PBS, homogenized, and processed for gDNA isolation as previously described (69). gDNA samples from individual animals (n = 5 per group) were pooled prior to analysis to generate a single representative sample per tissue and condition. Quantification of bacterial burden was carried out using a custom probe targeting the Cb com1 gene (Table S2). Cb GEs were normalized to host DNA using a commercially available mouse β-actin probe (BioRad assay ID dMmuCNS292036842). Each ddPCR reaction contained gDNA template, target-specific primers, and probes for both com1 and β-actin, with no-template controls included to monitor for contamination. Droplet generation was performed using the QXDx Automated Droplet Generator (Bio-Rad), which partitions each reaction into thousands of nanoliter-sized droplets. PCR amplification was subsequently carried out, and fluorescence signals were measured using the Qx200 Droplet Reader (Bio-Rad). Absolute quantification of target DNA was determined based on the fraction of positive droplets, and data were analyzed using QX Manager Standard Edition (Bio-Rad). Bacterial burden was expressed as the ratio of com1 to β-actin signal for each sample. Representative 1D and 2D droplet plots, along with sample annotations, are provided in the Supplementary Data File (<u>S12 Fig)</u>. Because gDNA samples from multiple animals were pooled prior to ddPCR analysis, each condition yielded a single composite measurement rather than independent biological replicates. All ddPCR Ssamples were prepared and analyzed at the Texas A&M Institute for Genome Sciences and Society (TIGSS). For measuring Cb burden in Fig 10, we used TaqMan quantitative PCR (TaqMan-qPCR) using primers for Cb CBU1910 (*com1*) as previously described (69). The sequence of the *com*1 TaqMan-qPCR primers and probe are listed in the S2 Table.

**Fig 10.**
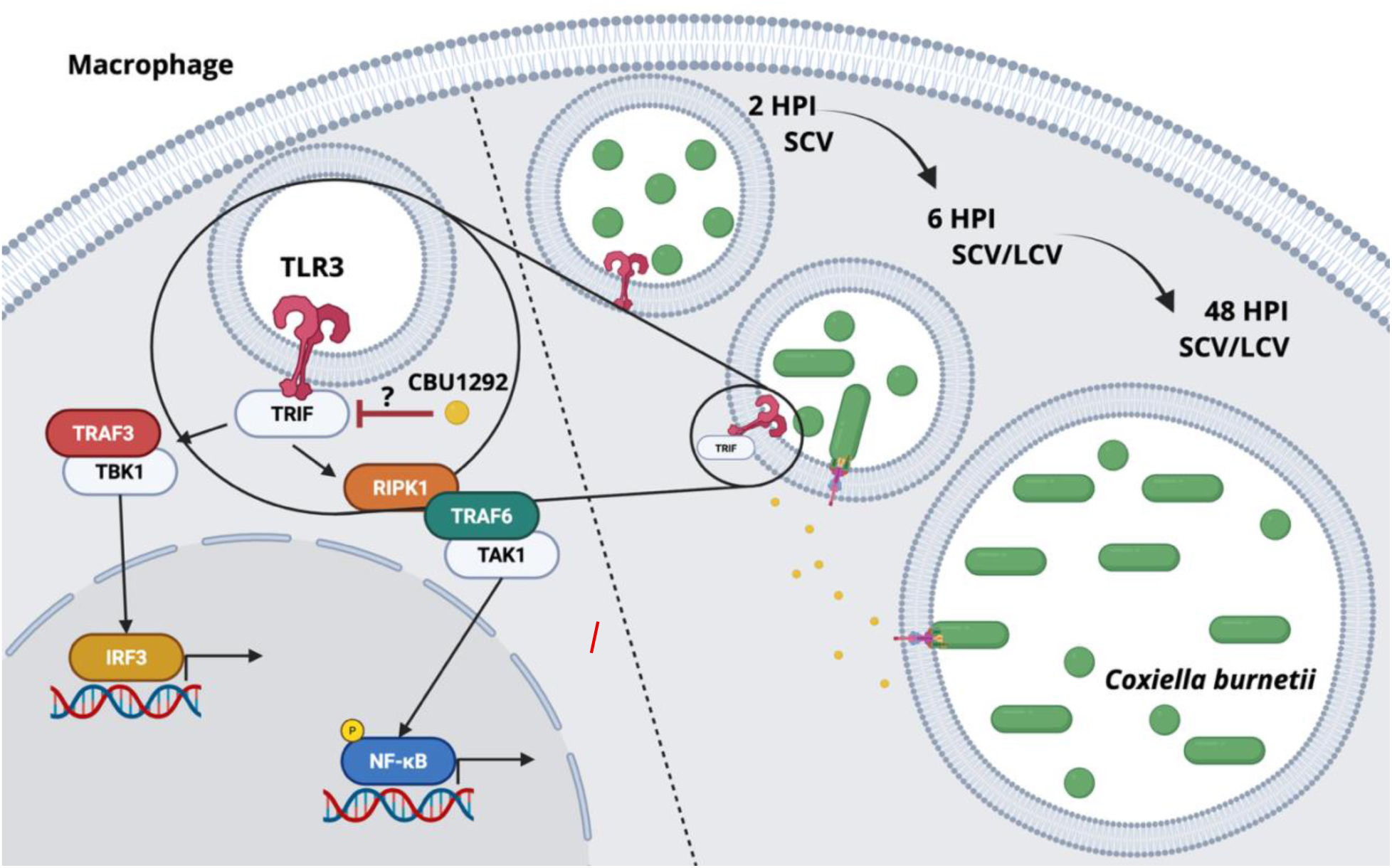
Conclusions and Working Model. In this study, we demonstrate that Cb suppresses host TLR3 signaling to dampen both NF-κB- and IRF3–dependent the transcriptional responses. This inhibition is T4SS-dependent and becomes apparent by ∼6 hpi *in vitro*. By 48 hpi, host TLR3 is no longer detected on mature CCVs formed by wild-type Cb. In contrast, TLR3 remains associated with CCVs formed by the T4SS-deficint DotA mutant, suggesting that Cb actively disrupts TLR3 recruitment through a T4SS-dependent mechanism mediated by an as-yet unknown effector. Furthermore, we show that Cb infection inhibits the association between TRIF and TRAF6 in a T4SS dependent manner, thereby blocking downstream NF-κB activation. Our data further suggest that multiple Cb T4SS effectors contribute to the modulation of host innate immune signaling, including the effector CBU1292 (AnkK), which appears to participate in regulating TLR3/TRIF-dependent responses downstream of TRIF as mutants are unable to inhibit NF-κB- and IRF3–dependent transcriptional responses. In addition, our observations are consistent with the involvement of distinct, as-yet unidentified effector(s) in the late-stage exclusion of TLR3 from the CCV.

### Immunofluorescence assays, Confocal microscopy, and Quantitative colocalization analysis

For time-course infections, 1 x10^6^ Hela cells were infected at an MOI of 25 with either wild-type NMII or T4SS-deficient DotA mutant in a 24-well tissue culture plate with coverslips. At 6 and 48 hpi, cells were fixed with 4% paraformaldehyde in 1x phosphate-buffered saline (PBS) for 15 min at room temperature (RT) and IFAs were performed. For immunolabeling, fixed cells were permeabilized and sequentially labeled with primary and secondary antibodies. All antibodies used in this assay are listed in the S1 Table. Briefly, NMII or DotA was labeled with guinea-pig anti-Coxiella antibody (red), while host TLR3 was stained with either rabbit anti-human TLR3 antibody (green). The host CD63 lysosomal marker was stained with mouse anti-human CD63 antibody (magenta), and Hoechst 33342 (blue) was used for DNA staining. Coverslips were mounted onto glass slides using Prolong Gold Antifade (Molecular Probes) and viewed by LSCM imaging using an Olympus Fluoview FV3000 confocal microscope. The quantitative colocalization analysis for NMII-TLR3 and DotA-TLR3 including the calculation of Pearson’s correlation coefficients, was conducted using the NIH ImageJ software and the JACOP plugin, following previously established methods (67). In three independent experiments, the 6hpi samples were analyzed by scoring three bacteria per infected host cell for Pearson’s coefficient colocalization in a total of 100 mammalian host cells. Similarly, for the 48hpi samples, one mature CCV was assessed in a total of 100 mammalian host cells for Pearson’s coefficient colocalization across three independent experiments. The authors acknowledge the assistance of the Integrated Microscopy and Imaging Laboratory at Texas A&M Health Science Center for these set of experiments and analysis.

### Co-Immunoprecipitation Assay for assessing host TRIF-TRAF6 interaction

Hela Cells were seeded overnight at 2 x 10^5^ cells and transfected with a Flag-tagged humanTRAF6 plasmid (listed in the S1 Table) using PolyJet (SignaGen). At 12 hours post transfection, cells were infected at an MOI of 25 with either wild-type NMII or T4SS-deficient DotA mutant for 12 hours. At 24 hours post transfection, infected cells were either mock treated or treated with 2 μg/ml of poly I:C (TLR3 agonist) for another 30 mins. Flag-tagged human TRAF6 co-immunoprecipitation (IP) assays were performed with anti-FLAG M2 affinity gel (Sigma-Aldrich) and following manufacturer’s recommendations. Briefly, cells were lysed for 1 hour on ice with 50mM Tris [pH 7.4], 150 mM NaCl, 1 mM EDTA, 0.075% NP-40, and protease inhibitors. Lysates were centrifuged for clearing the nuclei and cellular debris, and 10% of the Lysate was stored at - 80 degrees Celsius as input “whole cell lysate” (WCL). The remaining 90% of the cell lysate was incubated overnight with prewashed anti-FLAG M2 affinity gel. To minimize the denaturation of Flag M2-antibody, sample elution was achieved by using 2x Laemmli sample buffer lacking fresh β-mercaptoethanol (Bio-Rad). Proteins in the input WCL and output IP eluate were resolved by SDS-PAGE. Following western blotting, samples were probed with anti-FLAG, and anti-TRIF antibodies as listed in the S1 Table.

### TRIF-dependent NF-κB-inducible SEAP reporter assay to assess differential NF-κB activation by Cb NMII and DotA strains

For measuring NF-κB activation in the presence or absence of TRIF, THP-1-blue NF-κB-SEAP cells (TRIF Control) and TRIF knockout THP-1-NF-κB-SEAP cells (TRIF KO) from Invivogen were utilized (refer S1 Table). 1 × 10^6^ cells were seeded in 24-well tissue culture plates and infected with either wild-type NMII or T4SS-deficient DotA mutant at an MOI of 25. Cell culture supernatant from each sample was harvested at 12hpi, and NF-κB-inducible SEAP levels were determined using QUANTI-Blue Solution (Invivogen). Absorption was measured at 635nm with a PerkinElmer EnVision 2104 multilabel reader. For measuring TLR3-induced NF-κB-inducible SEAP activation in TRIF KO cells, either NMII or DotA (MOI 25) was added for 2 hours, coupled with poly I:C stimulation across a dosage gradient at 2hpi. Cell culture supernatant from each sample was harvested at 12hpi, and NF-κB-inducible SEAP levels determined as described above.

### Cb T4SS effector transposon mutant screen assay

To identify Cb T4SS effectors that suppress NF-κB activation, a NMII Himar1 transposon (Tn) mutant clone library with 32 NMII effector mutants was screened using THP1 NF-κB-Lucia reporter cells sourced from Invivogen (20). A total of 1 × 10^6^ THP1 NF-κB-Lucia cells were seeded in a 24 well tissue culture plate. Cells were either mock treated (negative control) or stimulated with 10^7^ heat-killed *Listeria monocytogenes* (positive control) or 200 ng/mL of *E. coli* LPS O111:B4 (positive control). The previously established CBU1639::Tn served as a positive control for NF-κB-Lucia induction (20). Test samples included wild-type NMII, the T4SS-deficient DotA mutant, and 31 NMII T4SS effector Tn mutants, which were added to THP1 NF-κB-Lucia cells at an MOI of 25. At 12 and 24 hpi, cell culture supernatants were collected, and NF-κB-inducible luminescence levels were measured in triplicate using the QUANTI-Luc coelenterazine-based luminescence detection assay (Invivogen). To measure Lucia bioluminescence, PerkinElmer EnVision 2104 multilabel reader was utilized. In total, three independent experiments were conducted, and the data obtained were normalized to those of the mock-treated control. This normalization allowed for the calculation of NF-κB fold induction. To assess whether CBU1292 contributes to suppression of IRF-dependent signaling, we utilized the CB*U1292*::Tn (*ankK*::Tn) mutant to measure IRF-Lucia reporter activation at 24 h post-infection (hpi). THP-1 IRF-Lucia reporter cells (InvivoGen) were seeded at a density of 1 × 10⁶ cells per well in 24-well tissue culture plates and allowed to equilibrate prior to infection. Cells were either mock-treated (negative control) or infected with wild-type NMII Cb or the CBU1292::Tn mutant under identical conditions. At 24 hpi, cell culture supernatants were collected, and IRF-inducible luciferase activity was quantified using the manufacturer’s recommended luminescence detection assay (InvivoGen). Luminescence measurements were performed in technical triplicate for each condition, and data were recorded as relative luminescence units (RLU) using the Cytation 5 plate reader (Agilent-Biotek).

### Complementation of the CBU1292 (*ank*K) Tn mutant in Cb NMII

The CBU1292 (*ank*K) open reading frame (ORF) was amplified using gene-specific primers for the wild-type NMII sequence and cloned into the pJB-proBA-2x-HA Coxiella expression plasmid (generously provided by Drs. Paul Beare and Bob Heinzen (NIAID Rocky Mountain Laboratories, Montana), using Gibson cloning. This plasmid was then used as a template to generate a complementation vector construct with Tn7 L and R sites flanking p1169-HA-CBU1292 and a p1169-kanamycin resistance marker for selection in Cb. To generate the complementation mutant, CBU1292::Tn (*ank*K::Tn) mutant strain was transformed with the Tn7-p1169-2xHA-CBU1292 complementation repair plasmid according to Dr. Paul Beare’s previously described protocol (70, 71). Following PCR confirmation of complementation in the population, cultures were plated on 1x ACCM-2-agarose plates and grown for 7 days to isolate individual colonies for gDNA extraction (70, 71). Individual colonies were scaled up and gDNA was isolated to confirm the presence of an additional copy of HA-tagged CBU1292 in the Cb genome (S11A and S11B Fig). All plasmids used for generating CBU1292::Tn (*ank*K::Tn) complementation (comp) strain are listed in the S1 Table. Additionally, all primers associated with this assay are listed in the S2 Table. For THP-1-Lucia NF-κB reporter cell infections with CBU1292::Tn (*ank*K::Tn) complemented strain, 1 x 10^5^ cells were infected with mCherry-expressing NMII (wild-type), DotA, 1292::Tn (*ank*K::Tn) mutant strain, or CBU1292::Tn (*ank*K::Tn) comp strain at MOI = 25. At 2 hpi, 20µg/ml of poly I:C was added to the media containing the infected cells. At 12 and 24 hpi, media was removed from the wells in triplicate for Luciferase assay. To measure NF-kB activity in response to infection, QUANTI-Luc^TM^ reagent (InvivoGen) was used according to manufacturer’s instruction. Relative luminescence was measured using the Cytation 5 plate reader (Agilent-Biotek).

### Statistical analysis / visualization and scientific illustration

For statistical analysis and visualization of experimental data, GraphPad Prism 10 was utilized. Additionally, Bio-Render was used for creating the scientific illustrations presented in Fig 11.

## Acknowledgements

We wish to thank Dr. Rahul Raghavan and Dr. Brandon E. Luedtke for the critical reading of this manuscript. This research was supported by National Institutes of Health grant NIH 5R01AI090142 and 5R21AI147123.

